# The glycine-arginine-rich motif of 53BP1 modulates RNA interactions necessary for its liquid-liquid phase separation during DNA Damage Response

**DOI:** 10.64898/2026.01.30.702603

**Authors:** Federica Terraneo, Marta Ceccon, Oscar Sapkota, Tongyin Zheng, Matteo Brizioli, Angela dello Stritto, Samara Cummings, Simone Codispoti, Giuseppe Ossolengo, Fabrizio Orsenigo, Serena Magni, Andrea Gottinger, Andrea Mattevi, Monika Fuxreiter, Nicolas L. Fawzi, Francisca Lottersberger, Fabio Giavazzi, Fabrizio d’Adda di Fagagna

## Abstract

The DNA damage response relies on the rapid assembly of repair factors into foci with properties of liquid-liquid phase separation, driven by *de novo* transcription of damage-induced RNAs. 53BP1 is a key component of these condensates, yet the molecular determinants driving this process remain unknown. Here, through computational, structural and *in vitro* approaches, we identify the oligomerization domain of 53BP1 and its glycine-arginine-rich (GAR) motif as crucial for RNA interactions and phase separation. Biophysical characterization reveals that 53BP1-RNA condensates can progressively mature into a more stable state, and that GAR mutants display aberrant material properties. Using a cellular model of telomere fusion events, we demonstrate that the GAR motif is essential for 53BP1-mediated DNA repair, which depends on the combined contributions of RNA binding and appropriate condensate biophysical properties. Therefore, RNA-driven 53BP1 condensation is functionally required to maintain genome integrity.

## INTRODUCTION

Cellular organization has traditionally been studied through the lens of membrane-bound organelles, which physically compartmentalize biochemical processes. However, recent advances in cell biology have revealed the existence of membraneless organelles, which can form through liquid-liquid phase separation (LLPS, also referred to as phase separation into liquid-like biomolecular condensates). In LLPS, weak, multivalent interactions among biomolecules (particularly nucleic acids and proteins containing intrinsically disordered regions (IDRs) and low-complexity domains (LCDs)) drive a homogeneous solution to demix into coexisting dense and dilute phases^1–3^. The resulting structures dynamically assemble and disassemble to compartmentalize and concentrate specific components, thereby promoting and regulating a wide range of cellular functions, with an important impact in both physiology and disease^4–7^. Notably, RNA molecules are frequently enriched in biomolecular condensates, acting as structural scaffolds and regulatory elements^8,9^.

A critical biological context in which LLPS plays a role is the DNA damage response (DDR), a coordinated set of events that implements detection, signaling, and repair of DNA lesions, safeguarding genome integrity against endogenous and exogenous insults^10,11^. DNA double-strand breaks (DSBs) represent the most severe form of damage due to their potential to trigger chromosomal rearrangements and loss of genetic information, and, if not properly repaired, cellular senescence and cell death fueling organismal aging and cancer^10–12^. While the DDR was originally considered exclusively as a set of protein-mediated events, mounting evidence points to a critical role for RNA, in particular its *de novo* synthesis at DSB^13^. We and others demonstrated that, following a DSB, the MRE11-RAD50-NBS1 complex (MRN) senses the lesion and recruits RNA polymerase II, which transcribes bidirectionally long non-coding RNAs (damage-induced long non-coding RNAs, dilncRNAs) from and toward the exposed DNA ends of DSB^14,15^. dilncRNAs are subsequently processed locally by the endoribonucleases DROSHA and DICER into shorter RNAs (DNA damage response RNAs, DDRNAs)^16,17^ that pair to complementary dilncRNAs and together promote the targeted recruitment of DDR factors at DSB in the form of DDR foci^14–16,18^.

Among key DDR foci components is 53BP1, a multidomain scaffold protein that regulates DSB repair pathway choice, promoting non-homologous end joining (NHEJ) by limiting DNA end resection and facilitating fill-in synthesis, alongside its role in damaged chromatin mobility and checkpoint activation through p53 signaling^19^. We and others previously discovered and reported that 53BP1 foci at DSB have liquid-like properties^20,21^, with RNA driving this process, providing an efficient mechanism for the rapid, on-demand, and reversible compartmentalization of the DDR machinery at DSB sites. Indeed, inhibition of synthesis or processing of damage-induced RNAs, their degradation or inhibition disrupts 53BP1 condensation and impairs DNA repair^20^. Together, these findings establish DDR foci as phase-separated compartments whose formation is critically dependent on local *de novo* transcription of damage-induced RNAs at DSBs. However, the mechanisms governing the formation and regulation of 53BP1 condensates by RNA remain poorly understood and whether this has a functional impact remains unknown. In this study, we identify the molecular determinants of 53BP1 responsible for its RNA-mediated phase separation in DDR. Combining computational analyses, *in vitro* RNA binding assays with recombinant 53BP1 fragments, phase separation assays and nuclear magnetic resonance (NMR) spectroscopy, we show that the oligomerization domain (OD) and its C-terminal disordered region containing a glycine-arginine-rich (GAR) motif are essential for RNA-driven LLPS. Beyond establishing their liquid-like nature, complementary measurements of material properties reveal a progressive maturation of 53BP1-RNA condensates toward a more stable configuration, characterized by increased surface tension and reduced molecular mobility while retaining liquid-like behavior. Finally, using a well-established cellular model of telomere dysfunction to recapitulate DSB repair, we demonstrate that the GAR motif is essential for 53BP1-mediated NHEJ, highlighting the functional dependency on RNA-driven 53BP1 condensation to support genome integrity.

## RESULTS

### Computational analysis of 53BP1 predicts extensive intrinsic disorder and fuzzy interactions critical for droplet formation

To gain an overall structural picture of human full-length 53BP1 and to identify regions potentially involved in phase separation, we first adopted a computational approach. 53BP1 has been characterized to bear a minimal region required for efficient DNA damage-induced focus formation, encompassing the OD, a GAR motif, a Tudor domain, and the ubiquitination-dependent recruitment (UDR) motif, which enable the interaction of 53BP1 with chromatin (histones γH2AX, H4K20me2 and H2AK15ub)^19,22^. Outside the minimal focus-forming region, the C-terminal tandem BRCT (BRCA1 C-terminal) domain is responsible for mediating 53BP1-γH2AX interaction and 53BP1-p53 interaction^19^. We used the IUPred2A method^23^ to predict the disorder propensity of 53BP1. The tool identified 53BP1 as a protein characterized by pronounced disorder, with long IDRs (**Fig. 1a**), a common trait among proteins undergoing LLPS through non-specific interactions, which allow molecules to sample a wide variety of different configurations^24^. Next, we explored whether these disordered regions retained their conformational heterogeneity upon interactions. FuzPred analysis^25^ indicated that full-length 53BP1 contains extensive regions with high probability of disordered binding (**Extended Data Fig. 1a**) that results in a heterogeneous ensemble of conformations. This suggests a strong propensity for dynamic multivalent contacts able to support phase separation.

**Fig. 1:**
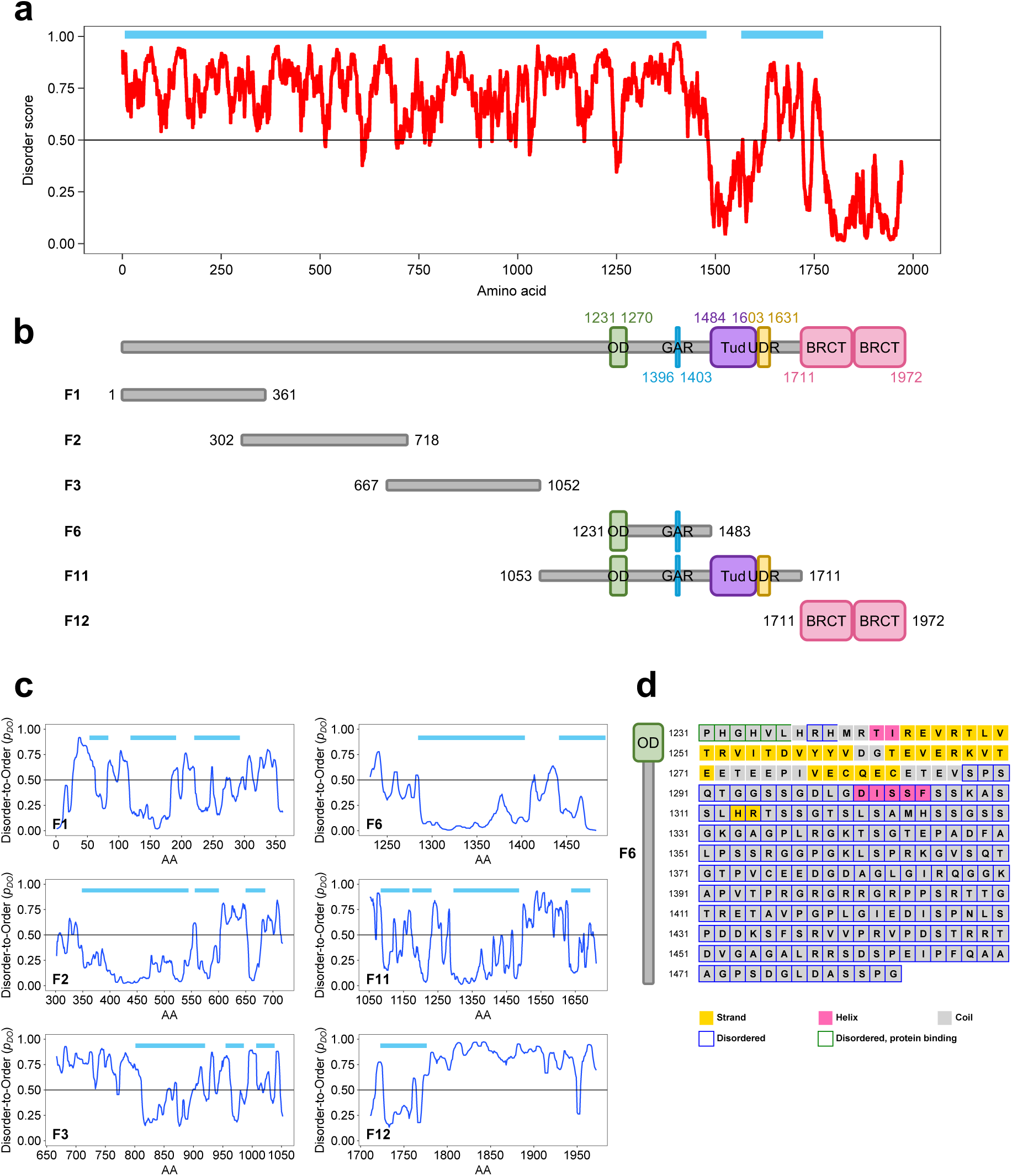
Computational analysis of 53BP1 predicts extensive intrinsic disorder and fuzzy interactions critical for droplet formation. **(a)** Graph plotting the predicted disorder score of human 53BP1 amino acid sequence, as calculated by IUPred2A. Scores above 0.5 (threshold marked by the horizontal black line) indicate predicted IDRs. Full-length protein was used as input. Horizontal blue lines mark regions of the protein which remain disordered upon binding (based on FuzPred predictions). **(b)** Schematic representation of full-length 53BP1 highlighting annotated protein domains (top), and the fragments designed for this study (bottom, from F1 to F12). OD = oligomerization domain, GAR = glycine-arginine-rich motif, Tud = Tudor domain, UDR = ubiquitination-dependent recruitment motif, BRCT = BRCA1 C-terminal domain. **(c)** Graphs plotting the probabilities of residues to undergo disorder-to-order transition upon binding (p_DO_) for individual 53BP1 fragments, as computed by FuzPred. Scores higher than 0.5 (threshold marked by the horizontal black line) indicate transitions to ordered states upon binding. Horizontal blue lines mark regions of the fragment which remain disordered upon binding. **(d)** Predicted secondary structure of the 53BP1 F6 fragment as computed by PSIPRED. Disordered residues are outlined in blue. Structured residues are colored in yellow (strand) and pink (helix).

To characterize which region of 53BP1 is responsible for RNA-mediated LLPS, we divided the full-length protein into individual consecutive, partially overlapping fragments encompassing its known structural and functional domains (**Fig. 1b**), based on insights from previous studies^21,26^. Using the same *in silico* computational approach, FuzPred predicted that F1, F2, F6 and F11 fragments of 53BP1 tend to retain their conformational heterogeneity and sample a diverse set of bound states, typical of the dynamic interactions required for liquidity (**Extended Data Fig. 1b**). Based on the FuzDrop method^27^ all investigated fragments contain one or multiple droplet-promoting regions (DPRs), which facilitate partitioning into condensates. In particular, F6 (comprising the OD followed by a short C-terminal disordered segment) and F11 (comprising OD, Tudor and UDR) have long DPRs (**Extended Data Fig. 1b**). The repetition of DPRs, such as in F1, F2, F3 and F11 also increase heterogeneity of the bound conformations. Given that the stability of the droplet state may stem from hydrophobic forces^24,27^, we screened the 53BP1 fragments for motifs that are embedded in disordered regions yet tend to undergo disorder-to-order transition (**Fig. 1c**). The presence of such motifs may indicate the ability to drive condensate formation. F3, and to lesser extent F1 and F11, contain segments with a local compositional bias, which drive them toward ordered structures upon assembly (**Fig. 1c**). In contrast, the F6 fragment has a long, 222-residue region lacking such features. In addition, we investigated the context-dependent interaction behavior of these segments, characterized by the multiplicity of binding modes (MBM), which can indicate the likelihood of converting the liquid-like droplet state into a more solid-like state^28^. Using the FuzDrop method we observed that the DPRs in all fragments exhibit context-dependent behaviors, except F6 (**Extended Data Fig. 1b**). This indicates that F6 tends to retain its heterogeneous state. This observation prompted a detail examination of the structural features of 53BP1 F6 fragment. Sequence-based PSIPRED prediction^29^ of 53BP1 F6 secondary structure highlighted its predominantly disordered nature, characterized by a low abundance of secondary elements such as helix and strand, except for the OD (**Fig. 1d**). We conclude that 53BP1 (and the F6 fragment in particular) has high LLPS potential, primarily driven by its intrinsic disorder and DPRs.

### 53BP1 OD and its C-terminal disordered region are sufficient for RNA-mediated LLPS in vitro

To experimentally validate our *in silico* predictions and to delve deeper into the contributions of different protein regions to RNA-driven LLPS, we subcloned, expressed and purified the 53BP1 fragments described above (**Fig. 1b**) as recombinant proteins in *Escherichia coli* (**Extended Data Fig. 2**), and conducted *in vitro* phase separation experiments. Under the conditions employed, none of the fragments could undergo phase separation on their own (**Fig. 2a**). The addition of PEG (polyethylene glycol), a synthetic polymer commonly used as crowding agent to promote phase separation *in vitro*, induced droplet formation for both F6 and F11 (**Extended Data Fig. 3a**), suggesting that both fragments possess the intrinsic capacity to undergo phase separation. However, in the presence of increasing amounts of total human cells RNA, only 53BP1 F6 fragment underwent phase separation (**Fig. 2a**) as quantified by the increasing number and size of F6 droplets detected by differential interference contrast (DIC) microscopy, consistent with RNA-mediated phase separation. These observations are in agreement with the previous computational results (**Fig. 1c**), which identified F6 as the most disordered, DPR-rich fragment with the highest tendency to form fuzzy interactions with low MBM, therefore the most prone to phase separate. Additionally, these results indicate that the disordered N-terminal portion of 53BP1 alone is not essential for RNA-mediated 53BP1 LLPS, consistent with a previous report^21^.

**Fig. 2:**
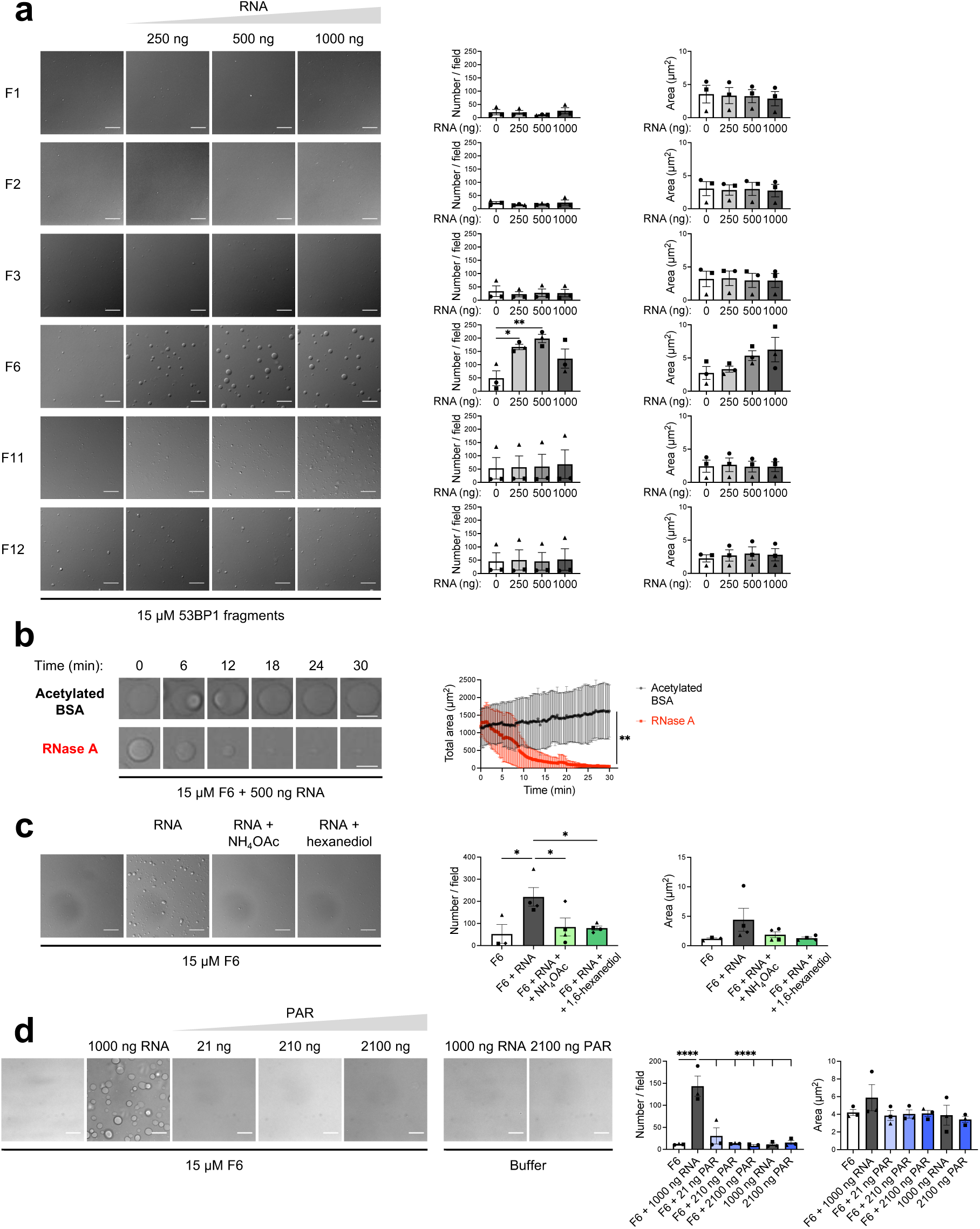
53BP1 OD and its C-terminal disordered region are sufficient for RNA-mediated LLPS *in vitro*. **(a)** Representative images of 53BP1 fragment droplets formation in presence of increasing amounts of RNA, with their relative quantification of droplets number per field and mean droplet area. Data are presented as mean ± SEM from *n* = 3 independent experiments. Each data point represents the mean of multiple fields from a single replicate. Statistical analysis was performed using one-way ANOVA, followed by Dunnett’s post hoc test against “0 ng RNA” group (* *P* < 0.05, ** *P* < 0.01). Differential Interference Contrast (DIC) channel is shown; scale bar = 10 µm. **(b)** Representative images of F6 droplets dissolution over time after RNase A treatment, with their relative quantification. RNA was added to the F6 fragment, droplets were allowed to form and then incubated with RNase A or acetylated BSA as a control. The graph shows the total droplets area over time after treatment addition. Data are presented as mean ± SD from *n* = 5 independent experiments. Statistical analysis was performed at the endpoint using an unpaired *t*-test (** *P* < 0.01). DIC channel is shown; scale bar = 5 µm. **(c)** Representative images of F6 droplets formation in presence of 1000 ng RNA and 100 mM ammonium acetate (NH_4_OAc) or 5% 1,6-hexanediol. Data are presented as mean ± SEM from *n* = 4 independent experiments. Each data point represents the mean of multiple fields from a single replicate. Statistical analysis was performed using one-way ANOVA, followed by Dunnett’s post hoc test against “RNA” group (* *P* < 0.05). DIC channel is shown; scale bar = 10 µm. **(d)** Representative images of F6 droplets formation in presence of increasing amounts of PAR, with their relative quantification of droplets number per field and mean droplet area. F6 with 1000 ng RNA was used as a positive control. Data are presented as mean ± SEM from *n* = 3 independent experiments. Each data point represents the mean of multiple fields from a single replicate. Statistical analysis was performed using one-way ANOVA, followed by Dunnett’s post hoc test against “1000 ng RNA” group (**** *P* < 0.0001). DIC channel is shown; scale bar = 10 µm.

To further confirm the requirement of RNA for 53BP1 F6 condensates formation and maintenance, we treated the droplets with RNase A to degrade RNA or with acetylated Bovine Serum Albumin (BSA) as a negative control. While with BSA the droplets’ area slightly increased over time (due to fusion events), upon RNase treatment F6 droplets completely dissolved within minutes (**Fig. 2b** and **Supplementary Video 1a,b**). Following RNase A inhibition by addition of RNaseOUT (a commercial reagent), re-addition of cellular RNA prompted condensates reformation (**Extended Data Fig. 3b** and **Supplementary Video 1c**), further demonstrating the RNA dependency of 53BP1 condensates and the reversibility of the process.

To prove the liquid-like nature of the F6 fragment droplets, we treated the condensates with ammonium acetate (NH_4_OAc), previously reported to dissolve RNA-mediated condensates^20,21,30^, and 1,6-hexanediol, used to disrupt weak hydrophobic interactions^20,31–33^. Both treatments hindered the formation of F6 droplets (**Fig. 2c**), pointing to the liquid-like nature of the condensates. Next, we wondered whether similarly negatively charged biopolymers could drive F6 phase separation similarly to RNA. To test this, we performed *in vitro* phase separation assays with poly(ADP-ribose) (PAR), which has features favorable to liquid demixing and has been observed enriched in some biomolecular condensates^34,35^, or with genomic DNA. Notably, F6 condensates failed to form in the presence of PAR (**Fig. 2d**), while genomic DNA induced the formation of solid aggregates rather than liquid-like droplets (**Extended Data Fig. 3c**), consistent with previous reports on the behavior of long DNA molecules^36,37^, altogether emphasizing the specificity of RNA as a driver of 53BP1 liquid-like assembly.

Altogether, these results demonstrate that the 53BP1 OD and its adjacent C-terminal disordered region (contained within the F6 fragment) are sufficient to drive RNA-mediated phase separation into liquid-like condensates *in vitro*.

### Disordered lysine/arginine-rich regions in 53BP1 F6 fragment mediate RNA interactions

To gain further insight into the structure of 53BP1 F6, we conducted NMR studies on the purified recombinant F6 fragment. The initial heteronuclear single quantum coherence (HSQC) spectrum showed backbone amide resonances with limited dispersion in the ^1^H dimension (**Fig. 3a**, left), consistent with the largely disordered nature of this fragment as suggested by computational analyses. We successfully assigned the majority of 53BP1 F6 residues in the 2D ^1^H-^15^N HSQC spectrum with the exception of residues in the OD and its immediate vicinity. We hypothesized that this was due to the OD forming large complexes that tumble slowly in solution, rendering these residues invisible in the NMR spectrum^38^. Consistent with this hypothesis, we observed a significant decrease in peak intensities for the assigned residues close to OD (**Fig. 3a**, top right). Additionally, we noticed another peak intensity dip around residue 180-220, suggesting that this region may also participate in 53BP1 self-interaction or undergo conformational exchange in solution. To gain insights into the secondary structure of 53BP1 F6, we conducted NMR secondary shifts and chemical shift-based secondary structure analysis using the SSP program^39^ (**Fig. 3a**, mid and bottom right). Here, positive values near 1 indicate α-helical characteristics and negative values near –1 suggest β-strands, while values near 0 are consistent with disorder. The results revealed that the NMR-visible segments of 53BP1 F6 are mostly disordered, with some transient secondary structure characteristics, validating experimentally our *in silico* predictions.

**Fig. 3:**
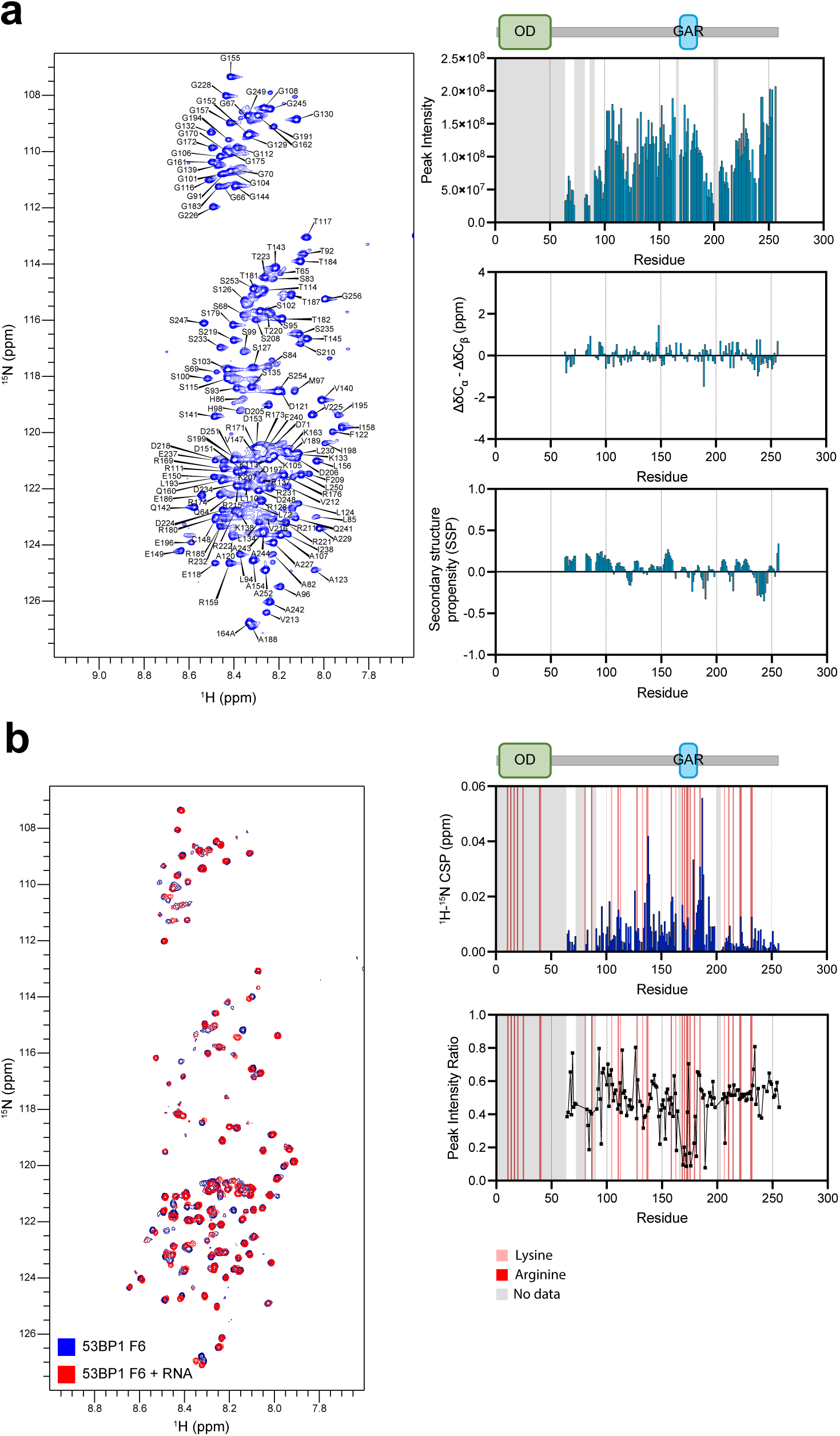
Disordered lysine/arginine-rich regions in 53BP1 F6 fragment mediate RNA interactions. **(a) (Left)** ^1^H-^15^N HSQC spectrum and backbone assignments of 53BP1 F6. **(Right)** ^1^H-^15^N HSQC resonance intensities for the assigned residue positions (top); the gray bars indicate residues with missing resonances. NMR experimental secondary chemical shifts (middle) and NMR chemical shift-based secondary structure analysis (bottom) using the SSP program. HSQC spectra were collected on *n* > 3 independent 53BP1 F6 samples. **(b) (Left)** ^1^H-^15^N HSQC overlay of 53BP1 F6 spectra with (red) and without (blue) the addition of Torula yeast RNA. **(Right)** Chemical shift perturbation (top) and peak intensity reduction (bottom) induced by the addition of Torula yeast RNA mapped to 53BP1 F6 sequence. Peak shifts observed upon adding RNA suggest nearby RNA interactions. Lysine and arginine residues are highlighted in pink and red, respectively; the gray bars indicate missing peaks. HSQC spectra were collected on *n* > 3 independent 53BP1 F6 samples.

To pinpoint the specific regions of the protein involved in RNA interaction, we recorded ^1^H-^15^N HSQC spectra using the F6 fragment in the presence or absence of total RNA extract (**Fig. 3b**, left). By comparing the two spectra, we mapped the induced chemical shift perturbations (CSP) to the 53BP1 F6 residues, revealing interactions within several crucial regions enriched in arginine and lysine (**Fig. 3b**, right). Notably, the most prominent CSPs and intensity reductions were observed around the GAR motif of 53BP1, located adjacent to its OD, implicating this region as a key mediator of protein-RNA interactions.

Collectively, these structural insights suggest that the combination of intrinsic disorder and RNA interactions at the GAR motif may provide the molecular basis for 53BP1 RNA-mediated phase separation.

### The glycine-arginine-rich (GAR) motif of 53BP1 is essential for RNA interactions and LLPS

RGG/GAR motifs, characterized by short arginine-glycine-rich repeats, are commonly found within IDRs, facilitating interactions between proteins and RNA molecules^40,41^, and are enriched in proteins that localize to biomolecular condensates^41^. 53BP1 GAR motif is highly conserved among orthologs (**Fig. 4a**), and was previously shown to have a role in mediating interaction with DNA^42,43^ and, more recently, RNA^44^. Although it was demonstrated that the unstructured sequence stretches C-terminal to the OD domain positively contribute to 53BP1 LLPS^21^, the specific involvement of the GAR motif in the RNA-dependent liquid demixing of 53BP1 remained unknown.

**Fig. 4:**
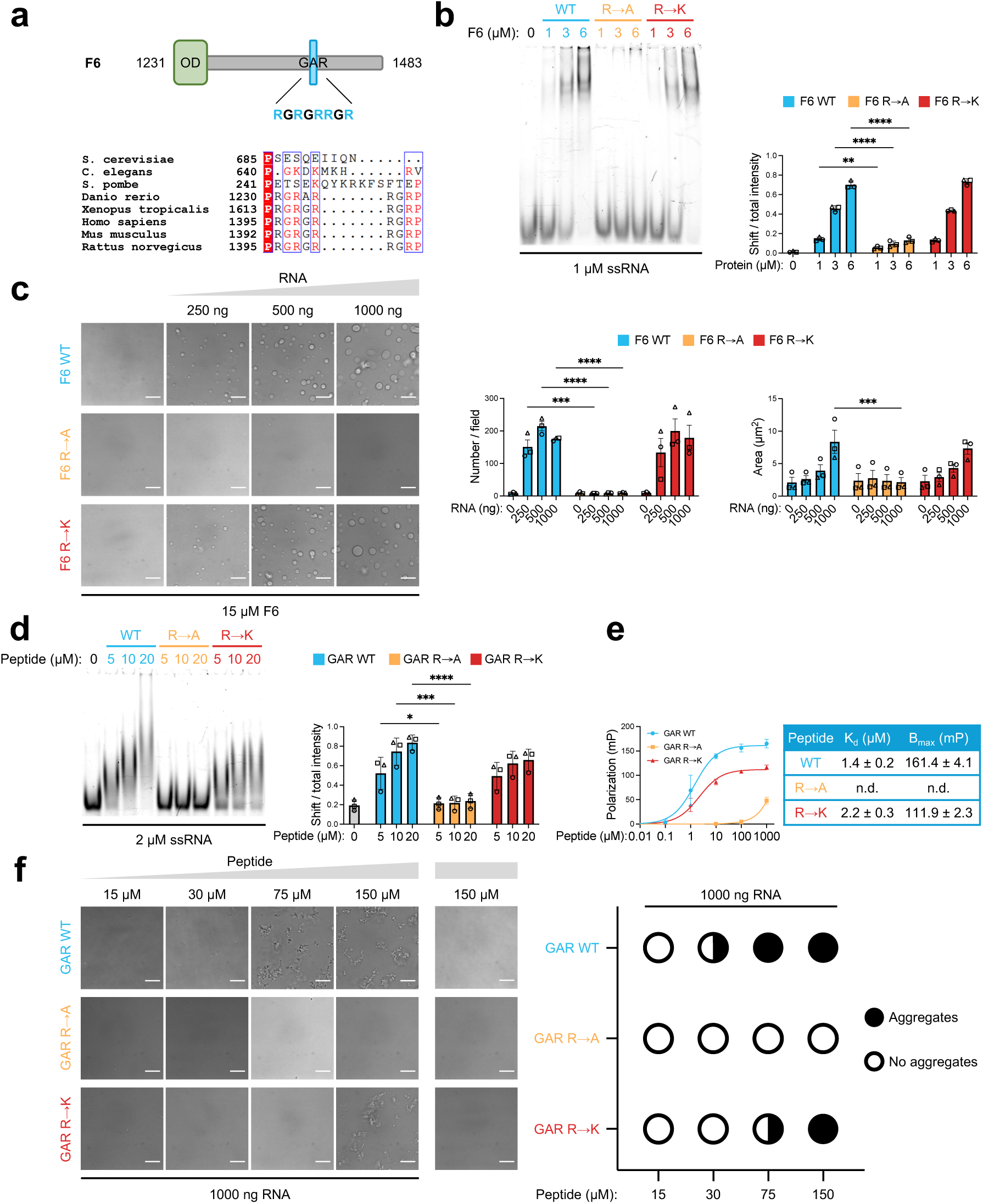
The GAR motif of 53BP1 is essential for protein-RNA interactions and LLPS *in vitro*. **(a) (Top)** Schematic representation of 53BP1 F6 fragment sequence, highlighting the OD and the GAR motif. **(Bottom)** Sequence alignment of the GAR motif region of 53BP1 orthologs from *Saccharomyces cerevisiae* (RAD9), *Caenorhabditis elegans* (hsr-9), *Schizosaccharomyces pombe* (crb2), *Danio rerio*, *Xenopus tropicalis*, *Homo sapiens*, *Mus musculus*, and *Rattus norvegicus*. **(b)** EMSA showing a titration of WT, R→A, R→K F6 fragments against a fixed concentration of 63-nt synthetic ssRNA (1 µM). Quantification was performed by calculating the ratio of bound RNA band intensity over total lane intensity for each condition. Data are presented as mean ± SD from *n* = 3 independent experiments. Statistical analysis was performed using two-way ANOVA, followed by Dunnett’s post hoc test against “F6 WT” group at each concentration (** *P* < 0.01, **** *P* < 0.0001). **(C)** Representative images of WT, R→A, R→K F6 droplets formation in presence of increasing amounts of RNA, with their relative quantification of droplets number per field and mean droplet area. Data are presented as mean ± SEM from *n* = 3 independent experiments. Each data point represents the mean of multiple fields from a single replicate. Statistical analysis was performed using two-way ANOVA, followed by Dunnett’s post hoc test against “F6 WT” group at each concentration (*** *P* < 0.001, **** *P* < 0.0001). DIC channel is shown; scale bar = 10 µm. **(d)** EMSA showing a titration of WT, R→K, R→A GAR peptides against a fixed concentration of 63-nt synthetic ssRNA (2 µM). Quantification was performed by calculating the ratio of bound RNA band intensity over total lane intensity for each condition. Data are presented as mean ± SD from *n* = 3 independent experiments. Statistical analysis was performed using two-way ANOVA, followed by Dunnett’s post hoc test against “GAR WT” group art each concentration (** *P* < 0.01, **** *P* < 0.0001). Two-way ANOVA revealed a significant main effect of the protein variant (**** *P* < 0.0001) and RNA concentration (* *P* < 0.05), but no significant interaction between the two factors, indicating that the effect of protein variant on droplet formation was consistent across all RNA concentrations tested. **(e)** Binding curves between 21-nt ssRNA and WT, R→A, R→K GAR peptides, obtained by FP. Polarization values were plotted against peptide concentration, while RNA concentration was kept constant (50 nM). Data are presented as mean ± SD from *n* = 4 independent experiments. The data points were fitted by non-linear regression using a one-site specific binding model. The solid line represents the best-fit curve. The apparent dissociation constants (K_d_) and maximum binding capacities (B_max_) are listed in the table (± SEM; n.d. = not determinable under tested conditions). **(f) (Left)** Phase separation assay showing a titration of WT, R→A, R→K GAR peptides against RNA (1000 ng). The experiment was performed 4 times, with similar results. DIC channel is shown; scale bar = 10 µm. **(Right)** Graph showing peptide concentrations categorized as positive (filled circle) or negative (open circle) for aggregate-like structures, with half-and-half circles indicating variability in the presence of aggregate structures at that peptide concentration.

Given the abundance of positively charged amino acids in the GAR motif of 53BP1, we hypothesized that its electrostatic interaction with RNA drives phase separation. To test this, we generated F6 mutants where all five arginine residues within the GAR motif of 53BP1 were replaced with alanine (F6 R→A, thus abolishing the motif’s positive charge) or lysine (F6 R→K, to discriminate a specific contribution of arginine residues beyond positive charge). To assess the RNA binding ability of F6 mutant fragments, we performed electrophoretic mobility shift assays (EMSA) using a synthetic 63 nucleotide single-stranded RNA (63-nt ssRNA) and increasing amounts of purified recombinant proteins. While F6 WT bound ssRNA in a concentration-dependent manner, F6 R→A mutant failed to interact with RNA, suggesting a role for the GAR motif in RNA interactions. Conversely, F6 R→K mutant exhibited no significant difference in RNA binding compared to the WT fragment (**Fig. 4b**). Next, we tested F6 mutants in *in vitro* phase separation assays with increasing amounts of RNA. Consistent with binding results, F6 R→K mutant retained the ability to form droplets as the F6 WT, while this process was completely abolished in the case of F6 R→A mutant (**Fig. 4c**), strongly supporting a phase separation mechanism facilitated by protein-RNA interactions mediated by the GAR motif.

To focus specifically on the isolated GAR motif, we synthesized 24-amino-acid-long peptides encompassing the GAR motif of 53BP1, both in its wild-type form (GAR WT peptide), alanine-mutated form (GAR R→A peptide), or lysine-mutated form (GAR R→K peptide) (**Extended Data Fig. 4a**). When performing EMSA with these peptides and the 63-nt ssRNA, GAR R→A peptide showed no interaction, while GAR R→K peptide showed comparable binding to the WT peptide (**Fig. 4d**), paralleling our findings with full F6 mutants. To independently validate these conclusions, we performed fluorescence polarization (FP), using increasing amounts of GAR peptides while maintaining a fixed concentration of a 6-FAM-labeled 21-nt ssRNA. The resulting binding curves revealed a complete abrogation of RNA binding in the case of the GAR R→A substitution, while the GAR R→K mutant showed an approximately 1.5-fold increase in the dissociation constant (K_d_) compared to the WT, indicating a slightly reduced binding affinity (**Fig. 4e**). Subsequent *in vitro* phase separation assays with these peptides in the presence of RNA demonstrated that the GAR WT peptide interacted with RNA, forming clearly detectable aggregate-like structures. We hypothesize that the peptides produced solid structures instead of liquid droplets because of their high charge density and lack of disordered, flexible regions flanking the GAR motif. In concordance with the binding assays, alanine substitutions within the GAR motif completely abolished the formation of these solid-like structures, while lysine substitutions altered the threshold required for aggregate formation at higher peptide concentrations compared to GAR WT peptide (**Fig. 4f**). Altogether, these results highlight an arginine-specific contribution to the RNA-53BP1 interaction process and phase separation.

With the recent release of AlphaFold 3, which enables AI-based structural prediction of proteins and their interactions with ligands and nucleic acids^45^, we performed structure predictions to complement the results from the binding assays with the GAR mutants. A single copy of the F6 WT, R→A and R→K sequences and copy of the 63-nt ssRNA sequence were used as input, and the predictions were run 10 times for each condition with 10 autogenerated seeds, to assess reproducibility. In the resulting predictions (**Extended Data Fig. 4b**), the GAR motif of F6 WT consistently localized in close proximity to the RNA, as confirmed by areas of reduced position error at the protein-RNA interface in the predicted aligned error (PAE) plot, consistent with putative interaction sites. In addition, the arginines within the GAR motif appeared to form multiple contacts with the RNA. In contrast, the mutated GAR motif in F6 R→A consistently positioned itself distant from the RNA, with no detectable interactions. The F6 R→K mutant exhibited a mixed behavior, in close proximity to the RNA in some predictions, but not in others generated from different random seeds. AlphaFold 3 predictions performed using single copies of GAR peptides yielded similar trends (**Extended Data Fig. 4c**). Although we are aware that AlphaFold’s performance with IDRs and RNAs is inherently limited, the consistent correlation between the predicted outcomes and the experimental data from binding assays (**Fig. 4b,d,e**) further supports a role for the GAR motif (and specifically of arginine residues) in RNA interactions.

Taken together, these data identify the GAR motif of 53BP1 as a critical driver of protein-RNA interactions and RNA-mediated LLPS, specifically relying on its arginine residues for this role. Importantly, our findings indicate that the positive charge of the motif *per se* makes an important contribution but may not be sufficient to explain its functions, as lysine mutants failed to fully recapitulate the phenotype of the wild-type protein.

### 53BP1 condensates undergo time-dependent maturation and mutation-dependent biophysical defects

To gain deeper insight into the material properties of F6 condensates we conducted an in-depth biophysical characterization. A hallmark of liquid-liquid interfaces is the presence of spontaneous surface fluctuations, commonly referred to as thermal capillary waves^46^. These interfacial fluctuations generate detectable intensity fluctuations in bright-field microscopy images (**Supplementary Video 2**), which we analyzed using Differential Dynamic Microscopy (DDM), a non-invasive and label-free technique that extract scattering-like information from time-lapse microscopy images^47,48^. For F6 droplets, we observed that the relaxation rate *Γ*(*q*) of these fluctuations exhibited a clean linear scaling *Γ*(*q*) ∼ *q* with the wavevector *q* (**Fig. 5a**). This linear scaling is an intrinsic signature of capillary waves at the interface between two viscous media, with *Γ*(*q*) = *v*_*DDM*_*q*, where *v*_*DDM*_ = 2*σ*/*η* is the propagation velocity of the capillary waves, determined by the ratio between surface tension *σ* and viscosity *η* of the more viscous phase^49^. Performing DDM analysis on the same F6 droplet over time, we discovered that the capillary velocity increased (**Fig. 5b**), along with decreased scattering amplitude (**Fig. 5b**, inset), stabilizing after ∼ 4 hours from droplet formation. When analyzing multiple F6 droplets, we observed a statistically significant difference in the capillary velocity between the day of the sample preparation and the subsequent days (**Fig. 5c**). These observations suggest that the system underwent equilibration over time. Specifically, the surface tension appeared to increase (as indicated by the decreased scattering amplitude, see also Online Methods), stabilizing the liquid-liquid interface of the droplets, whereas the viscosity did not exhibit any significant variation. This process likely reinforces droplet cohesion and structural integrity, sustaining 53BP1 foci and ensuring retention of DNA repair factors.

**Fig. 5:**
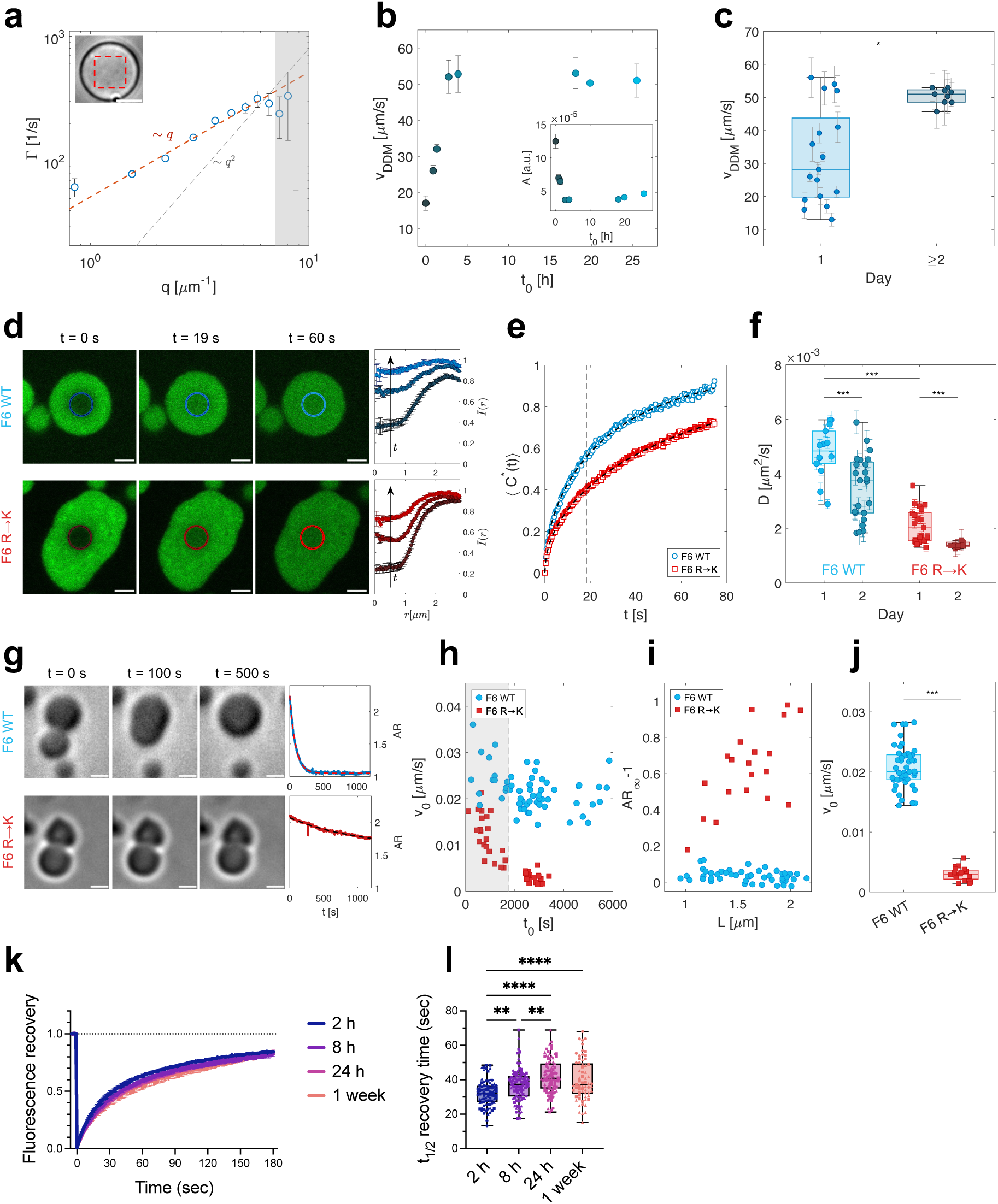
53BP1 condensates undergo time-dependent maturation and mutation-induced biophysical defects. **(a-c)** DDM analysis of F6 WT condensates. **(a)** Wavevector-dependent relaxation rate *Γ*(*q*) obtained from DDM analysis of a single F6 WT droplet. The dashed red line corresponds to the best fit to the rate with a linear model *Γ*(*q*) = *v*_*DDM*_*q*, from which *v*_*DDM*_ = (32 ± 1) *μm*/*s* is estimated. Top left: ROI where DDM is applied. The ROI is chosen to be smaller than the probed F6 WT droplet to avoid edge effects. Scale bar = 5 µm. **(b)** Time evolution of capillary velocity and amplitude (inset) computed from DDM for a single F6 WT droplet. **(c)** Capillary velocity for different F6 WT droplets estimated on the day of sample preparation and on subsequent days. Data are presented as box plots. Statistical analysis was performed using Mann-Whitney U-test or Wilcoxon rank sum test (* *P* < 0.05). **(d-f)** *In vitro* FRAP analysis of F6 WT and F6 R→K condensates. **(d)** FRAP measurements for two representative F6 WT and F6 R→K mutant condensates, respectively. Scale bar = 2 µm. Right: azimuthally averaged normalized intensity profiles measured at the time after photobleaching for which images are shown. **(e)** Normalized integrated time-dependent concentration after photobleaching ⟨*C*^∗^⟩(*t*) for F6 WT condensate (blue circles) and F6 R→K mutant condensate (red squares), respectively. Dashed lines correspond to the best-fitting curves based on the model described in Eq. (6). Vertical dashed lines indicate the time points reported in (d). **(f)** Effective diffusion coefficient *D* obtained by fitting 3D pure diffusion model with infinite boundary conditions to the normalized concentration for two different days after sample preparation. Data are presented as box plots. Statistical analysis was performed using Mann-Whitney U-test or Wilcoxon rank sum test (*** *P* < 0.001). **(g-j)** Coalescence analysis of F6 WT and F6 R→K condensates. **(g)** Representative images of F6 WT and F6 R→K mutant condensates undergoing coalescence in the stationary regime (*t* ∼ 30 min) during time after contact at *t* = 0. Scale bar = 2 µm. On the right the aspect ratio *AR* as a function of time. The dashed lines correspond to the best fit to the aspect ratio with a simple exponential model *AR*(*t*) = (*AR*_0_ − *AR*_∞_) exp(−*t*/*τ*) + *AR*_∞_. **(h)** Coalescence velocity *v*_0_ for different coalescence events observed in time after sample preparation for F6 WT condensate (blue circles) and F6 R→K mutant condensate (red squares), respectively. The white area marks the region *t*_0_ > 1.6 × 10^G^ *s* for which the sample has reached steady-state. **(i)** Asymptotic aspect ratio *AR*_∞_at steady-state as a function of the condensate radius *L*. **(j)** Coalescence velocity *v*_0_ for F6 WT (blue circles) and F6 R→K mutant (red squares) in the stationary state. Data are presented as box plots. Statistical analysis was performed using Mann-Whitney U-test or Wilcoxon rank sum test (*** *P* < 0.001). **(k-l)** FRAP analysis in irradiated BJ cells expressing a mCherry-tagged 53BP1 WT construct, performed at different time points after DNA damage. Cells were irradiated with 20 Gy, and FRAP was performed at 2 h, 8 h, 24 h, 1 week post-irradiation. **(k)** Fluorescence recovery time course after FRAP on BJ mCherry-53BP1 cells. Curves have been normalized to a 0 (intensity at the bleaching frame) to 1 (pre-bleaching intensity) scale for easier comparison. Data are presented as mean ± SEM from *n* = 3 independent experiments. **(l)** FRAP half-recovery times (*t_1/2_*) on BJ mCherry-53BP1 cells, derived from the curves shown in (k). Data are presented as box plots from 3 independent experiments (16-39 cells analyzed per time point per experiment), with different shapes indicating the different replicates. Statistical analysis was performed using one-way ANOVA, followed by Tukey’s post hoc test (** *P* < 0.01, **** *P* < 0.0001).

To investigate molecular mobility inside the droplets, we performed fluorescence recovery after photobleaching (FRAP) on F6 WT and F6 R→K samples doped with fluorescently labelled RNA (**Fig. 5d**). Assuming a 3D pure diffusion model with infinite boundary, we observed that the effective diffusion coefficient was significantly higher in the WT sample compared to the mutant (**Fig. 5f**), indicating lower internal mobility in the F6 R→K condensates and thus increased viscosity. Notably, a marked difference was observed between measurements taken on two consecutive days for both samples (**Fig. 5f**), suggesting that the system underwent aging with a gradual slowdown in molecular mobility. Importantly, fluorescence intensity fully recovered in both systems (**Fig. 5e**), indicating that, despite being slow, the sample retained liquid-like behavior. However, a 3D pure diffusion model with infinite boundary accounts only for diffusion while neglecting binding and reaction activity of the fluorescently labeled RNA within the sample, which may also play a role.

Another defining feature of LLPS is droplet coalescence, in which two liquid-like condensates fuse upon contact. We monitored coalescence during the first two hours following sample preparation (**Supplementary Video 3, Supplementary Video 4**). For both F6 WT and F6 R→K mutant condensates, the coalescence velocity *v*_0_ decreased over time, eventually setting on a constant value after *t* ∼ 30 min with a more pronounced decrease in F6 R→K mutant condensates (**Fig. 5h**). Strikingly, the fusions of the mutant droplets were arrested in the stationary regime (*t* ∼ 30 min), leading to aberrant shapes (**Fig. 5g**). While the asymptotic value of the aspect ratio *AR*_∞_in the stationary regime remained constant as a function of the condensate size *L* for F6 WT condensates, it exhibited a linear dependence on *L* for F6 R→K mutant condensates (**Fig. 5i**). Such a dependence is characteristic of condensates with a non-negligible elastic modulus, which counteracts surface tension and prevents the formation of perfectly spherical droplets upon fusion^50^. However, since passive coalescence occurring on a solid substrate is known to be strongly influenced by surface effects such as droplet pinning^51^, absolute physical properties (e.g., surface tension, viscosity, elastic modulus) cannot be straightforwardly quantified. Additionally, a significant difference was observed in the coalescence velocities evaluated in the stationary regime between F6 WT and F6 R→K mutant condensates (**Fig. 5j**).

Overall, the *in vitro* characterizations performed clearly demonstrate the liquid-like nature of F6 WT droplets, and an alteration in the biophysical properties of F6 R→K condensates. The slowed-down dynamics of F6 R→K droplets revealed by FRAP experiments (**Fig. 5d-f**) suggests either a higher viscosity of the dense phase or the presence of a non-negligible fraction of cross-linked molecular complexes. This latter scenario is consistent with the reduced fusion velocity of the mutant condensates observed in coalescence experiments (**Fig. 5g-j**), and with the linear dependence of the long-time aspect ratio *AR*_∞_ on the droplet size, which may indicate the presence of a significant elastic component^50^, possibly due to the formation of cross-linked structures inside the condensates. The notion that the main difference between F6 WT and F6 R→K mutant condensates relies in their bulk elastic properties is also compatible with the observation that their surface fluctuations, which are governed instead by interfacial properties, do not display significant differences (**Extended Data Fig. 5**).

To extend these *in vitro* observations *in vivo*, we exposed to X-rays normal human fibroblasts BJ cells expressing a 53BP1 construct fused to mCherry fluorescent protein, and we performed FRAP on fluorescent foci at different time points after irradiation. Fluorescence recovery was rapid at early time points, consistent with liquid-like behavior (**Fig. 5k**). Interestingly, we observed a progressive increase in the *t_1/2_* FRAP recovery time (**Fig. 5l**), indicating maturation into a more viscous, gel-like state, in agreement with our *in vitro* observations.

In summary, our comprehensive biophysical characterization revealed that 53BP1 F6 forms liquid-like condensates that undergo time-dependent maturation, both *in vitro* and in cells. Additionally, we have observed an aberrant *in vitro* LLPS behavior for the F6 R→K mutant, indicating that RNA binding *per se* is not sufficient to ensure WT-like biophysical properties, with potential impacts on DNA repair dynamics and efficiency.

### The GAR motif of 53BP1 is required for efficient NHEJ

One of the main functions of 53BP1 is promoting DNA repair of one-ended distal DSB by NHEJ. A convenient and well-established cellular system to recapitulate this phenomenon is the fusions of uncapped telomeres^52^. In this model, the removal of the shelterin protein TRF2 from telomeres generates free chromosome ends that are recognized as DSBs by the DDR, triggering fusions by NHEJ that are dependent on 53BP1 activity^52–54^. Therefore, to test the functional role of the GAR motif of 53BP1 in NHEJ, we complemented immortalized *Trf2^F/F^ 53bp1^-/-^* Mouse Embryonic Fibroblasts (MEFs)^54^ with either wild type human *53BP1* (*53BP1 WT*) or mutant alleles where all five arginine residues within the GAR motif were replaced with alanine (*53BP1 R→A*) or lysine (*53BP1 R→K*) (**Fig. 6a**).

**Fig. 6:**
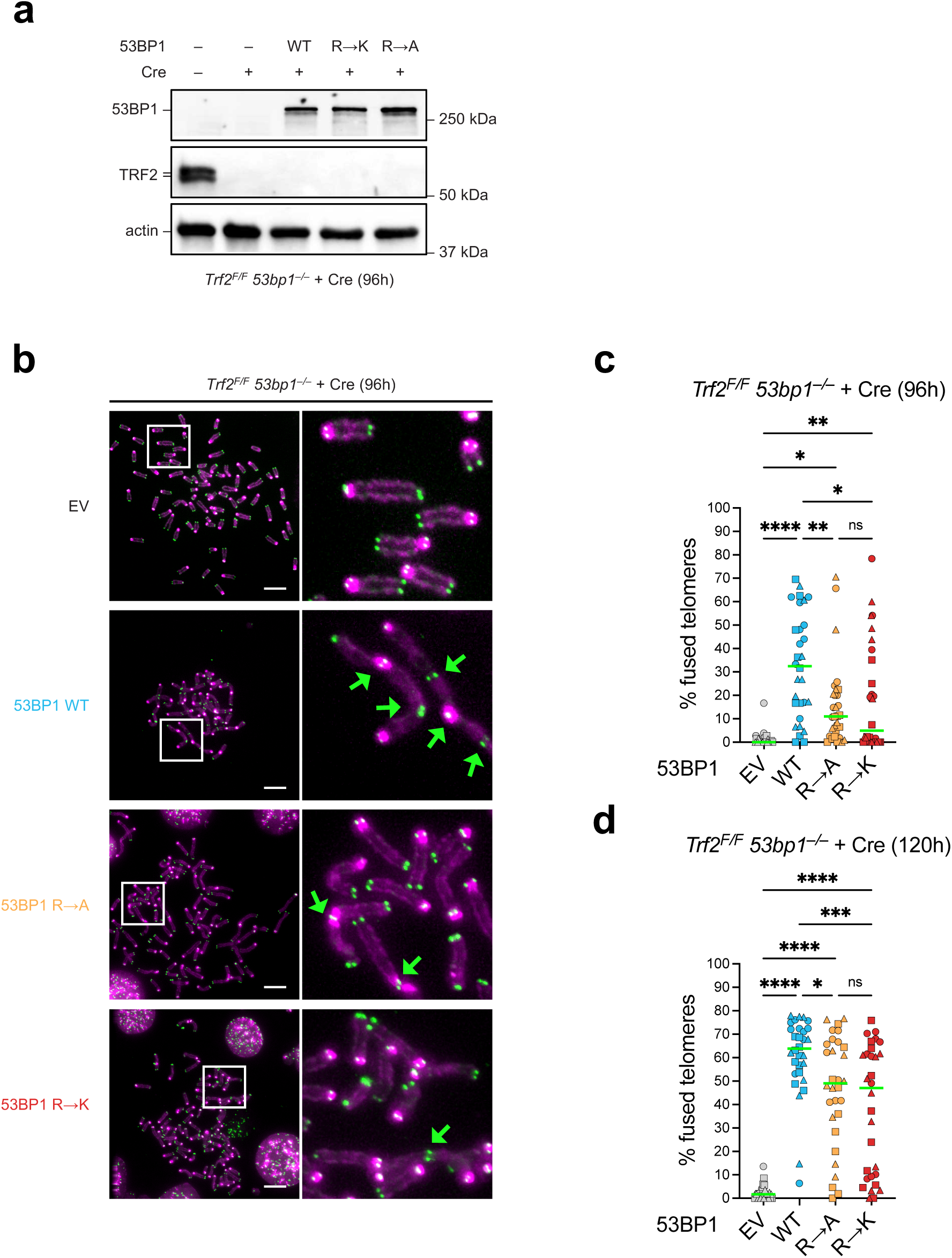
The GAR motif of 53BP1 is required for efficient NHEJ. **(a)** Immunoblot for 53BP1, TRF2 and actin (loading control) in *Trf2^F/F^ 53bp1^−/−^* MEFs expressing empty vector (EV), full-length wild-type 53BP1 (WT) or its GAR mutants (R→A or R→K), before or 96 h after Cre-mediated *Trf2* deletion with Hit&Run Cre. **(b)** Representative metaphase spreads in the indicated MEFs at 96 h after Cre-mediated *Trf2* deletion. Telomeres were detected with Alexa Fluor 488-OO-(TTAGGG)_3_ (green), DNA with DAPI (magenta). Arrows indicate telomere fusions. The boxed regions are enlarged in the right row. Scale bar = 10 µm. **(c-d)** Quantification of telomere fusions shown in (b) at 96 h **(c)** or 120 h **(d)** after Cre-mediated *Trf2* deletion. Data are presented as median from 3 independent experiments (10 metaphases each). Each data point represents the percentage of telomeres fused in one metaphase, with different shapes indicating the different replicates. Statistical analysis was performed using one-way ANOVA, followed by Tukey’s post hoc test (* *P* < 0.05, ** *P* < 0.01, *** *P* < 0.001, **** *P* < 0.0001).

As expected, *53bp1^-/-^* cells transduced with the empty vector control showed a minimal level of fusions after *Trf2* deletion with Hit and Run Cre recombinase^55^, consistent with the absence of 53BP1. MEFs complementation with *53BP1 WT* resulted in a time-dependent accumulation of interchromosomal fusions, with the median of telomeres engaged in fusion increasing from about 35% 96 h after *Trf2* deletion to about 65% after 120 h (**Fig. 6b-d**). Remarkably, both 53BP1 R→A and 53BP1 R→K mutants exhibited a strong reduction in the percentage of fused telomeres (about 10% and 5% at 96 h after *Trf2* deletion, **Fig. 6b,c**), indicating compromised function of 53BP1. Interestingly, GAR mutations seemed to affect the fusion kinetics, which in turn impacted the repair efficiency, rather than resulting in a complete functional defect: in fact, at a later time point (120 h) the fusion levels increased for all constructs, although the mutants still showed significantly lower fusion rates compared to WT (**Fig. 6d**). This observation aligns with the biophysical profile of the *in vitro* mutant droplets, which remain liquid-like but exhibit slowed-down internal dynamics and a non-negligible elastic component; such altered material properties likely hinder the optimal rate of telomere fusion without fully compromising the protein’s functional potential over longer timescales.

Altogether, these results indicate that alterations in the GAR motif impair the ability of 53BP1 to promote NHEJ, and therefore proper 53BP1 functions in *in vivo* repair depend on a critical combination of RNA-binding affinity and finely tuned biophysical properties of the condensates.

## DISCUSSION

DSBs represent a severe threat to genomic integrity, and defects in their repair are a primary driver of genomic instability and tumorigenesis^11,12^. As DDR has recently emerged to rely on LLPS for its spatial and temporal organization, elucidating the mechanisms that govern the formation of liquid compartments during this process is essential for identifying deregulated components that could be selectively targeted in pathological conditions. We and others have previously established that the DDR protein 53BP1 accumulates at sites of DNA damage in liquid-like foci^20,21^, dependent on site-specific RNA transcripts^20^. However, the precise mechanism remained largely unknown. In this study we dissected the molecular features governing this phenomenon, identifying 53BP1 OD and the adjacent C-terminal disordered region, specifically the GAR motif, as central drivers of protein-RNA interactions and phase separation, with GAR mutations affecting both RNA binding and the condensates material properties. Furthermore, we revealed that these condensates are not static, but undergo a progressive maturation toward a less dynamic state. Crucially, using a telomere-dysfunction model, we demonstrate that the specific arginine residues required for this RNA-mediated phase separation are also indispensable for efficient NHEJ in cells, thus establishing a direct functional link between the biophysical properties of 53BP1 condensates and genome integrity. Our results indicate that the disordered N-terminal portion of 53BP1 alone is not essential for RNA-mediated 53BP1 LLPS, which is consistent with a previous report^21^. Interestingly, while previous studies suggested the Tudor domain was essential for RNA interaction^14,56^, our data indicate it is not required for RNA-driven LLPS *in vitro*. This may be explained by the role of the Tudor domain in the recognition of damaged chromatin, mediating the interaction of 53BP1 with the histone mark H4K20me2 surrounding DSBs^57^ – therefore, *in vivo*, the absence of the Tudor domain may prevent the recruitment and localization of 53BP1 to the site of damage^56^ where nascent RNAs are transcribed, thereby hampering RNA binding and liquid demixing. Indeed, a more recent study reported that the recombinant Tudor domain of 53BP1 lacks the ability to bind RNA *in vitro*^58^. Taken together, these observations suggest a coordinated mechanism in which histone modifications act as initial recruiters, defining the chromatin domains where 53BP1 accumulates, while DSB-induced RNAs may subsequently function as local condensers, promoting 53BP1 phase separation at sites of damage.

Interestingly, we observed that the F11 fragment, which contains the F6 sequence and the Tudor domain, underwent PEG-induced but not RNA-induced LLPS. This intriguing observation may suggest that the Tudor domain in the context of the F11 fragment forms intramolecular interactions with the GAR motif, due to its ability to bind RG-rich peptides with low affinity^59^, thereby forming an auto-inhibitory conformation that may mask RNA access. Alternatively, the Tudor domain’s net negative charge may discourage recruitment of RNA and hence prevent condensation. These processes may both be at work and these hypotheses will need further investigations. In addition, our computational analysis highlights that the F6 fragment uniquely retains its conformational heterogeneity upon binding; conversely, the context-dependent behavior of the F11 fragment may favor conformational stabilization upon RNA interaction, selectively suppressing RNA-driven LLPS while preserving crowding-assisted condensation.

A central discovery of this work is the specific requirement for arginine residues within the GAR motif for functional RNA binding and condensate dynamics, beyond just positive charge. Mutagenesis experiments revealed that substituting arginines with lysines significantly reduced diffusivity and impaired fusion kinetics of 53BP1 condensates. This difference could be explained by the chemical versatility of the arginine guanidinium group, capable of forming complex hydrogen bond networks and cation-π interactions^41,60^, thereby promoting multivalency and sustaining condensates liquidity. Replacing arginine with lysine reduces both the affinity and the multivalency of RNA binding, shifting the system toward a kinetically trapped, gel-like state, demonstrating that distinct condensate material properties can arise despite comparable RNA binding and phase separation capacity. These observations align with previous studies showing that mutations or post-translational modifications (reported also in the RGG motifs) often lead to aberrant phase separation^6^. More broadly, both the sequence composition and amino acid patterning of RNA-binding proteins strongly influence their propensity to undergo LLPS and determine the material properties of the resulting condensates^61,62^. For instance, in FUS family proteins arginine has been shown to promote LLPS more effectively than lysine, possibly due to its ability to engage in interactions with aromatic residues via additional interaction modes^61,63^. Moreover, arginine-containing mixtures tend to favor liquid-like phase separation with aromatic residues, much more than lysine- or alanine-containing systems^64^. Considering the presence of aromatic rings in nucleobases, this may underlie both the morphological differences and altered dynamics observed in F6 R→K mutant droplets^65,66^.

Importantly, these biophysical alterations directly impact cellular functions. Using TRF2-deficient MEFs to trigger telomere uncapping and mimic a DSB, we demonstrated that both R→A and R→K mutations significantly impair 53BP1-mediated NHEJ, an effect that could be explained only by a combination of RNA binding and biophysical properties. Previous studies reported that (smaller) mutations or the deletion of the GAR motif of 53BP1 did not strongly affect NHEJ of TRF2-depleted telomeres^53,67^. However, since the defect was mild, it was not explored further. In addition, prior work also reported that deletion of the GAR motif or substitution of the arginines with alanines did not abolish (but reduced in some settings) 53BP1 localization to damage foci^42,56^, but without evaluating the repair outcome. In our experiments, mutating all five arginines of the GAR motif and investigating the dynamic over time allowed us to detect a significant reduction and delay in the fusions of telomeres upon TRF2 depletion. The observation that R→A (which cannot form condensates) and R→K mutants (which can still form condensates but with compromised fluidity *in vitro*) fail to support efficient telomere fusion to a similar extent suggests that RNA binding *per se* is necessary but not sufficient for proper function, and that condensates’ liquid state is essential for the rapid ligation of DNA ends *in vivo*. Interestingly, from a functional standpoint, a solidified or kinetically trapped assembly appears to be as detrimental as the total absence of phase separation. Together, these results establish a connection between a molecular-level alteration (an amino acid substitution), a biophysical defect (loss of condensate fluidity) and, ultimately, a compromised functional phenotype at the cellular level (reduced DNA repair). Therefore, proper condensate formation is not merely a matter of forming a phase-separated state in the context of DSB repair, but also of maintaining the correct material and dynamic properties necessary for downstream function, and efficient DNA repair occurs when RNA binding and condensate biophysical properties are both properly tuned.

We characterized the rheological evolution of these condensates, observing a time-dependent shift in their material properties, both *in vitro* and in live cells. This “aging” process suggests that persistent foci may naturally transition into more stable structures. Persistent foci are localized at genomic loci that are resistant to repair, like dysfunctional telomeres^68^ and in senescent cells^69^. While increased viscosity may protect irreparable DSBs from unwanted interactions with other nuclear components, the emergence of solid-like characteristics could impede resolution and sustain DDR signaling, contributing to maintenance of cellular senescence^68,70,71^. Such maturation is analogous to what has been observed in other RNA-binding protein (RBP) condensates, where liquid-like droplets can transition into viscoelastic gels or solids^72–74^. This is of great biological relevance, as solidification (often mediated by pathological aggregation and fibril formation) is a hallmark of neurodegenerative diseases^75–77^. For example, FUS and other prion-like proteins can form gels and aggregates inside droplets over time^78–80^, and disease-associated mutations can expedite these changes in the material state of the condensates^78,79^. Similarly, cancer-associated mutations can accelerate aberrant phase transitions^81^. Whether naturally occurring somatic mutations of 53BP1 influence condensate material properties and predispose to disease remains an intriguing avenue for future research. It is known that the MBM drives sampling of both disordered and ordered conformations, and cellular factors may trigger the shift in between these states: this switch may associate with binding mode promiscuity leading to cellular toxicity^82^. Our preliminary computational data (data not shown) indicate that certain cancer-associated 53BP1 mutations, which increase the susceptibility to changing of binding mode upon cellular cues (i.e. increase the MBM) as compared to the WT, are located within the F6 fragment region. Interestingly, mutations affecting the droplet-promoting probability (p_DP_) are notably scarce in this region. This may suggest that most mutations occurring within the F6 fragment sequence have the potential to alter the material properties of the droplets. Further investigations are warranted to test this hypothesis.

Furthermore, since arginines in the 53BP1 GAR motif can be methylated^42,43,83^, and this post-translational modification can modulate RNA binding affinity and LLPS in other RBPs^84–86^, future work may determine if the methylation status of the GAR motif changes during the DDR and how it influences phase separation.

Similarly, recent studies have reported emerging functions for the epitranscriptome in DDR pathways^87–90^, LLPS^91–93^ and RNA:DNA hybrids^89,94–97^. Considering the role of non-coding RNAs in 53BP1 recruitment and LLPS^14,20^, it is possible that RNA modifications may influence these processes, thereby offering new avenues for research.

In conclusion, this study redefines the GAR motif as a crucial driver of RNA-mediated 53BP1 condensation. We demonstrate that the specific physicochemical properties of arginine residues are essential not just for RNA interaction, but for maintaining the dynamic liquid state of the condensates. By linking the material properties of these foci to the efficiency of NHEJ, we provide evidence that the biophysical state of the repair compartment is a critical determinant of genome maintenance. These findings deepen our understanding of the interplay between protein disorder, RNA, and phase separation in the DDR, offering new perspectives on how dysregulation of these mechanisms may contribute to genomic instability and disease, and paving the way for new investigations and potential therapeutic approaches. Indeed, condensates are gaining increasing attention as therapeutic targets in neurodegeneration, infectious disease and cancer^98^, with the first condensate-modifying drug entering clinical trials for the treatment of Wnt-driven tumors. In addition, evidence shows partitioning of some anti-cancer drugs within liquid condensates, leading to their disruption^99^.

As an example of therapeutic use of agents that modulate LLPS, we previously reported that antisense oligonucleotides (ASOs) against telomeric dilncRNAs and DDRNAs dissolve DDR foci and ameliorate disease phenotypes in models of aging^100,101^ and neurodegeneration^102^, and can elicit a selective vulnerability in a specific class of cancers that exploit alternative lengthening of telomeres (ALT) for their survival^103^. This emphasizes the potential of targeting condensates as a valuable approach for therapeutic intervention across various pathological contexts.

## ONLINE METHODS

### Computational analysis of protein structure

Identification of IDRs of 53BP1 was carried out using IUPred2A^23^. Human 53BP1 protein sequence was used as input in the web interface to get a score between 0 and 1 for each residue, indicating the probability of a residue to be part of a disordered region. Values above 0.5 indicate a high probability of that residue residing in an unstructured region.

The binding modes of 53BP1 fragments, including identification of motifs undergoing disorder to order transition, were characterized by the FuzPred method. FuzPred predicts the context-dependent binding behavior of proteins, by distinguishing between ordered binding (when disorder-to-order transitions occur upon binding) and disordered binding (when disorder is largely retained in the complex)^25,104^.

Droplet-promoting regions were predicted using the FuzDrop method, which estimates the propensity of proteins to undergo LLPS^27,28^.

53BP1 F6 predicted secondary structure was computed using PSIPRED^29^.

For structure prediction using AlphaFold 3^45^, sequences of the F6 fragments or GAR peptides (in WT or R→A or R→K mutant forms) were uploaded to the AlphaFold 3 Server, together with 63-nt ssRNA, in a 1:1 ratio. Out of the 5 predictions resulting from each run, only the top ranked prediction was considered. This procedure was repeated 10 times with 10 auto-generated seeds. Predictions were processed using ChimeraX software^105^.

### Plasmids and cloning

For the expression of 53BP1 fragments as recombinant proteins, 53BP1 fragments were amplified from the plasmid encoding FL 53BP1 (pcDNA5-FRT/TO-eGFP-53BP1, 60813, Addgene) using different primer pairs [FF1/Rev361 for 53BP1 F1 (residues 1-361); FF302/Rev718 for 53BP1 F2 (residues 302-718); FF667/Rev1052 for 53BP1 F3 (residues 667-1052); FF1231/Rev1483 for 53BP1 F6 (residues 1231-1483); FF1053/Rev1711 for 53BP1 F11 (residues 1053-1711); FF1711/Rev1972 for 53BP1 F12 (residues 1711-1972)] and Phusion DNA polymerase (NEB). The inserts were ligated into the pGEX-6P-1 vector plasmid (kind gift of A. De Antoni, V. Costanzo’s lab, IFOM) within BamHI/XhoI restriction sites using the Quick Ligase kit (NEB). The final products were verified by DNA sequencing.

F6 fragment bearing R→A mutations in the GAR motif of 53BP1 (R1396A, R1398A, R1400A, R1401A, R1403A) was generated from pcDNA5-FRT/TO-eGFP-53BP1 (60813, Addgene) by running two PCR reactions using the primer pairs FF R(GAR)A/Rev1483 (for Fw fragment) and FF1231/Rev R(GAR)A (for Rev fragment). A third PCR was then run using the previously obtained Fw and Rev fragments and the primer pair FF1231/Rev1483 to obtain F6 fragment bearing R(GAR)A mutations (F6 R→A). The insert was ligated into the pGEX-6P-1 vector within BamHI/XhoI restriction sites using the Quick Ligase kit (NEB). The product was verified by DNA sequencing.

F6 fragment bearing R→K mutations in the GAR motif of 53BP1 (R1396K, R1398K, R1400K, R1401K, R1403K) was generated by PCR using the primer pair FF1231/Rev1483 on a pcDNA5-FRT/TO-eGFP-53BP1 R(GAR)K plasmid (mutated version of Addgene #60813, already in use in the lab, see next paragraph in this section). The obtained fragment was ligated into the vector pGEX-6P-1 within BamHI/XhoI restriction sites using the Quick Ligase kit (NEB) to obtain F6 fragment bearing R(GAR)K mutations (F6 R→K). The product was verified by DNA sequencing.

For the generation of pcDNA5-FRT/TO-eGFP-53BP1 R(GAR)K plasmid, R→K mutations (R1396K, R1398K, R1400K, R1401K, R1403L) in the GAR motif of 53BP1 were inserted by mutagenesis PCR on the plasmid pcDNA5-FRT/TO-eGFP-53BP1 (60813, Addgene) using the primers pair FF FL_R(GAR)K/Rev FL_R(GAR)K and Phusion DNA polymerase (NEB). The PCR product was digested with DpnI and ligated using the Quick Ligase kit (NEB). The final plasmid was verified by DNA sequencing.

For the telomere fusions experiment, 53BP1 full-length mutant bearing R→A mutations (R1396A, R1398A, R1400A, R1401A, R1403A) in the GAR motif of the protein was generated by mutagenesis PCR on the plasmid N-Myc-53BP1 WT pLPC-Puro (19836, Addgene) using the phosphorylated primers pair FF FL_R(GAR)A/Rev FL_R(GAR)A and Phusion DNA polymerase (NEB). The PCR product was digested with DpnI and ligated using the Quick Ligase kit (NEB). The final plasmid was verified by DNA sequencing.

53BP1 full-length mutant bearing R→K mutations (R1396K, R1398K, R1400K, R1401K, R1403L) in the GAR motif of the protein was generated by cut and paste from pcDNA5-FRT/TO-eGFP-53BP1 R(GAR)K (mutated version of Addgene #60813, already in use in the lab, see previous paragraph in this section) into N-Myc-53BP1 WT pLPC-Puro plasmid (19836, Addgene). Both plasmids were digested with AgeI/BlpI restriction enzymes, and the reaction products were run on a 1% agarose gel. Desired bands (backbone from pLPC plasmid and insert from pcDNA5 plasmid) were then excised from gel, extracted and ligated using the Quick Ligase kit (NEB). The final plasmid was verified by DNA sequencing.

### Expression and purification of recombinant GST-tagged PreScission protease

pGEX vector encoding N-terminal Glutathione S-Transferase (GST)-tagged 3C protease (PreScission protease) was a kind gift of Ario de Marco (Nova Gorica University). The construct was expressed in *E. coli* BL21 Star (DE3) pRARE2 in ZYM-5052 autoinduction medium for 16 h at 20°C. Harvested cells were resuspended in lysis buffer (50 mM Tris-HCl pH 8.0, 300 mM NaCl, 10% glycerol, 0.2% NP-40, 0.02% 1-thioglycerol, 1 mg/mL lysozyme, 2 μg/mL DNase I) and disrupted by sonication. The clarified supernatant was applied onto a GSTrap column on an ÄKTA Purifier (Cytiva) and washed extensively with wash buffer (50 mM Tris-HCl pH 8.0, 300 mM NaCl, 10% glycerol, 0.01% 1-thioglycerol). Bound protein was eluted with elution buffer (50 mM Tris-HCl pH 8.0, 300 mM NaCl, 10 mM reduced glutathione, 10% glycerol, 0.01% 1-thioglycerol); fractions were pooled together and dialyzed overnight against 1 L storage buffer (50 mM Tris-HCl pH 8.0, 150 mM NaCl, 10 mM EDTA, 20% glycerol, 0.01% 1-thioglycerol) at 4°C.

### Expression and purification of recombinant 53BP1 fragments

53BP1 fragments were expressed in *E. coli* BL21-CodonPlus (DE3)-RP. Cells were grown in Luria-Bertani (LB) medium until optimal density (OD_600_ = 0.6-0.8), and protein production was induced with 300 µM isopropyl-β-D-1-thiogalactopyranoside (IPTG) for 16 h at 17 °C. Harvested cells were resuspended in lysis buffer (50 mM Tris HCl pH 7.4, 50 mM NaCl, 5% glycerol, 2 mM DTT, 0.1% Triton X-100, 1 mM PMSF) supplemented with protease inhibitors (04693116001, Roche), stirred for 1 h at 4 °C, in presence of benzonase nuclease (E1014, Merck) and lysozyme, and then disrupted by sonication. The clarified supernatant was applied to a Pierce Glutathione Agarose resin (16100, Thermo Fisher) and washed extensively with high-salt (500 mM NaCl) and low-salt (50 mM NaCl) wash buffer. On-column cleavage of the GST tag was performed overnight at 4°C using PreScission protease. The flowthrough (containing the cleaved protein) was applied to a Resource Q chromatography column (17117901, Cytiva) pre-equilibrated in IEX buffer A (50 mM Tris pH 8.4, 50 mM NaCl, 5% glycerol, 1 mM DTT). The protein was eluted using a salt gradient (up to 1 M NaCl), concentrated and then applied to a Superdex 200 Increase 10/300 GL (28990944, Cytiva) pre-equilibrated in SEC buffer [50 mM Tris HCl pH 7.4 (pH 8.4 for F6 fragment), 150 mM NaCl, 5% glycerol, 1 mM DTT]. Protein-containing fractions were concentrated and stored in SEC buffer.

Screening of protein-containing fractions was performed using 4-12% Bis-Tris Acrylamide gels (Invitrogen) run in MES buffer (Invitrogen), stained with InstantBlue (Expedeon).

53BP1 F6 fragments bearing R→A or R→K mutations within the GAR region were purified following the same protocol as other 53BP1 fragments.

### 53BP1 F6 fragment expression and purification for NMR studies

The sequence coding for 53BP1 F6 was cloned into the pTHMT vector^106^ that encodes an N-terminal histidine-tagged maltose binding protein (MBP) followed by a TEV protease cleavage site and then the insert to make His-MBP-53BP1 F6, and subsequently transformed into BL21 Star (DE3) cells. For isotopically labeled protein, M9 minimal medium supplemented with appropriate isotopes was utilized. Cultures were grown at 37°C with continuous shaking until reaching an optical density at 600 nm (OD_600_) of 0.6-0.8. Expression of the fusion protein was induced by adding 1 mM Isopropyl β-D-1-thiogalactopyranoside (IPTG) and incubating the cultures at 37°C for 4 h. Following induction, the bacterial cells were harvested and lysed in the lysis buffer (20 mM NaPhos pH 7.4, 1 M NaCl, 10 mM Imidazole, 1 mM DTT) using a sonicator to disrupt the cell membranes. The lysed cell suspension was centrifuged, and the resulting supernatant containing the His-MBP-53BP1 F6 protein was collected. To purify the protein from cell lysate, a 5 ml HisTrap HP column was employed. The sample-loaded HisTrap column was washed with 20 mM NaPhos pH 7.4, 1 M NaCl, 10 mM Imidazole, 1 mM DTT buffer, then the bound protein was eluted with 20 mM NaPhos pH 7.4, 1 M NaCl, 300 mM Imidazole, 1 mM DTT buffer. The His-MBP tag was removed by first incubating the HisTrap-purified protein with TEV protease and then separating using a Superdex 200 (26/600) size-exclusion column, yielding a highly purified protein sample ready for further analysis.

### Mass photometry

Data were acquired using a TwoMP Instrument (Refeyn) equipped with an anti-vibrational table (Refeyn) using droplet-dilution mode by diluting 53BP1 F6 to 200 nM in a 20 μL drop of SEC Buffer. Data were fitted with a calibration obtained combining measurements of 50 nM Beta-amylase from sweet potato (BAM, Sigma Aldrich) and of 50 nM Bovine Serum Albumin (BSA, Sigma Aldrich). Data analysis was carried out using DiscoverMP v2.5 Software (Refeyn).

### Thermal unfolding

Purified 53BP1 F6 was diluted in SEC buffer at a final concentration of 2 mg/mL. Thermal unfolding was measured using Prometheus NT.48 Series High Sensitivity Capillaries (NanoTemper GmBH) and a Prometheus Panta NT.48 Instrument (NanoTemper GmBH) following nano Differential Scanning Fluorimetry (nanoDSF), backreflection, Static Light Scattering (SLS) and Dynamic Light Scattering (DLS) in a 25 to 95 °C temperature gradient with a ramp of 0.5 °C/min. Data analysis was carried out with PR.PantaAnalysis v1.8 Software by calculating the first derivative of the curves fitted by non-linear regression of the data.

### In vitro phase separation assay

A µ-Slide 18 well flat chamber (81826, Ibidi) was coated with a lipid layer as previously described^107^. Briefly, 10 mg/mL of a 1,2-Dioleoyl-*sn*-glycero-3-phosphocholine (DOPC, P6354, Sigma) solution in chloroform was dried and resuspended in LLPS buffer (25 mM Hepes pH 7.5, 150 mM NaCl, 1 mM DTT), then used to coat the chamber (1 h incubation, then washes with LLPS buffer). Purified 53BP1 recombinant fragments or synthetic 53BP1 GAR peptides (Mimotopes) were diluted in protein SEC buffer, mixed with PEG, total RNA extract, Poly(ADP-ribose) (PAR) polymer (Bio-Techne), or genomic DNA extract at the stated concentration, and incubated for 30 min at room temperature before microscopy analysis. When not specified, the final protein concentration was 15 µM, while the final RNA concentration was 40 ng/µL. When used, NH_4_OAc (100 mM) and 1,6-hexanediol (5%) treatments were added in the mixture as indicated. For experiments in which droplets were dissolved with RNase A, 100 ng of RNase A (Thermo Fisher) or 100 ng of acetylated BSA (Invitrogen) were added 30 min after droplets formation (dropping the reagent on the surface of the well). Immediately after treatment addition, a timelapse video was acquired every 10 sec for a total of 30 min. The mixture was then incubated with 40 U RNaseOUT (ThermoFisher) for 15 min to inactivate RNase A, followed by addition of extra 500 ng total RNA and imaging 30 min later. Images and timelapses were acquired using a DeltaVision Elite system (GE Healthcare) equipped with a IX71 microscope (Olympus) and a sCMOS camera and driven by softWoRx v.7.0.0. A UPlanSApo 60X/1.42 NA objective was used.

### Quantification of phase-separated droplets and measurement of shape parameters

In order to detect and quantify the droplets for each frame, a Napari (napari contributors 2019. doi:10.5281/zenodo.3555620) plugin to execute the nucleAIzer^108^ segmentation algorithm and its model “General – nuclei” was used. The resulting labelled images were analyzed with a customized Fiji^109^ script as follows. To detect each label as a droplet, the script converts the image in a binary mask setting a threshold and applying the watershed tool (https://imagej.net/imaging/watershed). The script uses the “Analyze particles” plugin to count the droplets and extract their shape descriptors (https://imagej.net/ij/docs/menus/analyze.html#ap): area, perimeter, minor and major axis, Feret diameter, circularity (4π*area/perimeter^2) and aspect ratio (major_axis/minor_axis).

### NMR sample preparation and NMR spectroscopy

Isotopically labeled protein samples of 53BP1 F6 were prepared in a buffer containing 20 mM NaPhos pH 6.7, 150 mM NaCl, 1 mM DTT, and 5% D_2_O. NMR spectra were recorded on Bruker Avance 600 MHz or 850 MHz ^1^H Larmor frequency spectrometers with HCN TCl z-gradient cryoprobes. Two-dimensional ^1^H-^15^N HSQCs were acquired using spectral widths of10.5 ppm and 30.0 ppm in the direct and indirect dimensions, with 3072 and 400 total points and acquisition time of 172 ms and 77 ms, respectively. Triple resonance experiments, including HNCACB, CBCA(CO)NH, HNCO, HNCA, and HNN, were conducted to establish sequential and side-chain resonance assignments. Acquisition times for the triple resonance experiments were 185 ms in the direct ^1^H dimensions, 24 ms in the indirect ^15^N dimensions, 15 ms in the indirect C’ dimensions and 6 ms in the indirect C_α_/ C_β_ dimensions. Spectral widths were 13 ppm in ^1^H, 22 ppm in ^15^N, 7.5 ppm in C’ dimension and 54 ppm in C_α_/C_β_ dimension. When stated, Torula yeast RNA was added to the sample prior NMR measurement. Data processing and analyses followed standard protocols using NMRPipe and CCPNMR Analysis 2.5 software^110,111^. 53BP1 F6 HSQC spectra were collected from more than three independent samples yielding similar results.

### Electrophoretic mobility shift assay (EMSA)

For EMSAs with 53BP1 F6 recombinant fragments, a 6% polyacrylamide native gel was cast and pre-run at 90V at 4°C with cold 0.5X TBE buffer. Purified 53BP1 F6 fragments, either WT or R→A and R→K mutants, were mixed with 3’-biotinylated 63 nt-long ssRNA (Dharmacon, as previously used^91^) in F6 SEC buffer. The reaction mixture was incubated at room temperature for 30 min, supplemented with 1:5 of 50% glycerol and loaded onto the gel. Gels were run with 0.5X TBE buffer at 120 V for ∼2 h, stained in SYBR Safe (S33102, Invitrogen) for 30 min, and de-stained with 0.5X TBE buffer at room temperature. Results from the gel were imaged using ChemiDoc MP (Bio-Rad) and quantified using Fiji^109^ by calculating the ratio of bound RNA band intensity over total lane intensity. For EMSAs with GAR peptides (Mimotopes), a 9% polyacrylamide gel was used instead.

### Fluorescence Polarization (FP) assay

For FP binding assay, 50 nM of FAM-labeled CXCR4 oligo in 1x siRNA buffer (Horizon Discovery) were mixed with increasing concentrations (from 0 µM to 1000 µM) of 53BP1 GAR peptides (Mimotopes) in water. Each measurement was performed in duplicate on a Tecan 200 plate reader in black 384-well microplates (3820, Corning) with 20 µL of mixture per well. The FP signals were recorded at room temperature using 485-nm excitation and 535-nm emission filters. The FP values were plotted against the log of the peptide concentrations, and the dissociation constant (K_d_) and the maximum binding capacity (B_max_) were obtained from fitting the points with nonlinear regression “One site – Specific binding” curve, as analyzed in GraphPad Prism 10.

### Differential dynamic microscopy (DDM) of interfacial fluctuations

The relaxation dynamics of thermal fluctuations occurring at the interface between the condensed and the dilute phase in phase-separating mixtures were measured using DDM. Samples were prepared as described in the *“In vitro liquid-liquid phase separation assay”* section above. Immediately after preparation, a drop of sample (volume 10 µL) was sandwiched between two glass coverslips separated by a silicone gasket of thickness 1.0 mm with a circular aperture of diameter 9 mm (GraceBio-Labs FastWells^TM^ reagent barriers). The inner surfaces of the glass coverslips were coated with BSA 3% w/w using the following protocol: the BSA drop (10 µL) was left for 20 min on the glass surface, then removed and two successive washing applying RNase free water for 10 min were performed; the surface was then let drying.

The sample was then let equilibrate for about 2 h at room temperature after the cell was sealed. Observations of the phase-separated droplets deposited on the bottom surface of the cell were performed at room temperature using an inverted microscope (Nikon Eclipse T*i*) equipped with a fast CMOS camera (Hamamatsu Orca Flash 4) under coherent bright field illumination and using a 60x (NA 0.70) objective.

An experiment consists in the acquisition of multiple (typically 3) image sequences (each one consisting of 20000 frames, captured at 800 fps) for each droplet. Many different droplets (typically N = 10) were considered in each experiment. To ensure suitable spectral resolution, droplets with radius larger than 2.5 µm were selected. Under these conditions, a square region of interest (ROI) of at least 40×40 pixels completely included within the droplet boundaries can be obtained, corresponding to a linear size of about 4.3 µm. Each collected image stack *I*(***x***, *t*) was analyzed according to the DDM processing scheme, leading the azimuthally averaged image structure function (ISF) *d*(*q*, Δ*t*) = ⟨x*Î*(***q***, *t* + Δ*t*) − *Î*(***q***, *t*|^2^⟩, where *Î* (***q***, *t*) is the spatial Fourier transform of *I*(***x***, *t*) and the symbol ⟨·⟩ indicates an average performed over the initial time *t* and the two-dimensional wavevectors ***q*** such that |***q***| = *q* (as in ^47,48^).

The following model

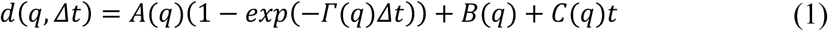

was then fitted to the ISF, leading to an estimate for the scattering amplitude *A*(*q*), the relaxation rate *Γ*(*q*), the noise floor *B*(*q*) and the drift term *C*(*q*). As discussed in ^112^, the last term in the above equation accounts for the presence spurious contributions due to rigid displacements, mainly due in the present case to mechanical vibrations and imperfect pinning of droplet on the surface,

Over the accessible *q*-range, roughly corresponding to interval [1, 10] *μm*^−1^, the obtained *q*-dependent relaxation rate invariably displays a clean linear dependence on *q Γ*(*q*) ≃ *v*_*DDM*_*q*. This peculiar linear dispersion relation clearly indicates that the origin of the optical signal cannot be attributed to diffusive motion (for which a quadratic dependence on *q* of the relaxation rate would be expected), but it is due instead to thermally exited capillary waves at the interface between the two phases (^49^ and Brizioli et al., in preparation). In fact, for wavevectors *q* much larger than the inverse of the so-called capillary length 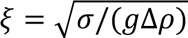, where *σ* is the interfacial tension between the two fluids, Δ*ρ* is their mass density difference and *g* is the acceleration of gravity, the relaxation rate of interfacial fluctuations is given by *Γ*(*q*) = (*σ*/2η)*q*, where *η* is an effective viscosity which, if two liquid phases have markedly different viscosities, corresponds to the viscosity of the condensed phase.

This enables identifying the coefficient *v*_*DMM*_, which we estimate by fitting a linear function to the experimentally obtained *Γ*(*q*), as the speed *v*_*DDM*_ = *σ*/2*η* of the capillary waves. The above-described procedure thus enabled obtaining, for each droplet, an estimate of the ratio *σ*/*η*.

The scattering amplitude *A*(*q*) in Eq. (1) is directly proportional to the power spectrum *S*(*q*) of the interfacial height fluctuations

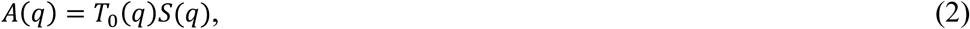

where the proportionality term *T*_0_(*q*) depends on the details of the imaging setup and on the refractive indexes of the two phases (^48^ and Brizioli et al. in preparation). For thermal capillary waves, if *q* ≫ *ξ*^−1^, *S*(*q*) is given by

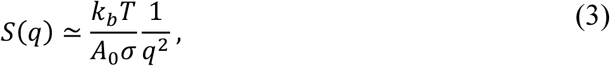

where *k_b_* is the Boltzmann constant, *T* is the absolute temperature, and *A*_0_is the area of the considered ROI (^46^ and Brizioli et al. in preparation). While possible in principle, a fully quantitative determination of *σ* from the scattering amplitude is challenging in practice, due to the undetermined proportionality term *T*_0_(*q*). However, if we consider a single droplet under constant observation conditions, any observed change in the measured average scattering amplitude *Ā* ≡ ⟨*A*(*q*)⟩_*q*∈[1.5,5]*μm*_^−1^ can be attributed to variation of the interfacial tension between the two fluids. In particular, under the above-described conditions

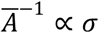

Monitoring the time evolution of *Ā*^−1^ thus enables tracking the overtime *relative* change of *σ* for an individual droplet.

### In vitro fluorescence recovery after photobleaching (FRAP)

The viscous properties and the molecular mobility within the condensate phase are also investigated by performing FRAP measurements. Samples are prepared according to the procedure outlined in section *“In vitro liquid-liquid phase separation assay”* with the addition of 1 µM ssCXCR4-FAM fluorescent RNA, and then sealed within cells as described in section *“Differential dynamic microscopy (DDM) of interfacial fluctuations”.* FRAP has been performed exploiting a laser scanning Confocal Microscope (Leica Stellaris 8) equipped with a 63x (NA=1.40) oil immersion objective (HC PL APO CS2) and a sCMOS camera Leica DFC9000 GTC. The imaging system is operated using the LASX (LEICA) Software. Photobleaching is achieved by using a 495 nm laser at maximum output for two cycles, each lasting approximately ∼ 470 ms. For a given droplet, bleaching is performed on a circular ROI of 2 µm-diameter. For each bleached ROI, a sequence of 5 frames (495 nm wavelength, output power=5%) at a frame rate of 2.7 frame per second is recorded prior bleaching. Immediately after bleaching, a sequence of 200 frames is captured at the same frame rate over a total duration of ∼ 75 s. Images cover an area of 23.1 *μm*^2^ (512×512 pixels). To monitor photobleaching during the acquisition, only droplets close to at least one other droplet, chosen as a reference, are considered.

For each FRAP measurement the pre- and post-bleaching sequences are analyzed. Pre-bleaching images are initially processed by convolving them with a Gaussian kernel of size 2 pixels to reduce noise. Subsequently, a threshold, selecting only the pixels with intensities above the 70^th^ percentile of the image intensity distribution, is applied. In the case of circular droplets, an automated algorithm for circle recognition is exploited to accurately identify the droplet boundaries and centroids. For non-circular droplets, a user-assisted algorithm is used to interactively identify the largest circle inscribed within a droplet. Since not all the droplets are pinned to the surface but they might jiggle close to it, a modified version of Image-J plugin *Stack-Reg* for rigid image registration is executed on cropped images centered on the droplet centroids and enclosing them. This modified plugin is applied to the image sequence obtained as combination of the pre- and post-bleaching subsequences, providing the transformation matrices of the rigid-body registration (translation and rotation) between pairs of images. Once the matrices are known, the centroid positions can be corrected over time, enabling accurate tracking of the droplets without relying on any user-dependent or automated thresholding, which may vary during acquisition due to photobleaching.

The normalized integrated time-dependent concentration after photobleaching can be defined as^113^

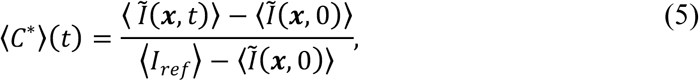

where the average ⟨·⟩ is taken over the bleached ROI. The term *Ĩ*(*x*, *t*) represents the corrected image intensity distribution at time *t*, which is defined as *Ĩ*(*x*, *t*) = *ξ*(*t*) · *I*(*x*, *t*), where *I*(*x*, *t*) is the captured image intensity distribution, and *ξ*(*t*) is a correction factor accounting for photobleaching during the acquisition. The correction factor *ξ*(*t*) is determined by monitoring the intensity of nearby, non-bleached reference droplets within the field of view and it is computed as 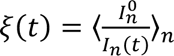 where *I* (*t*) is the intensity of a *n*-th reference droplet at time t, nd *I_n_*^0^ Is the pre-bleaching intensity of the same droplet. Finally, ⟨*I_ref_*⟩ represents the pre-bleaching intensity distribution of the bleached ROI.

The time evolution of ⟨*C*^∗^⟩(*t*) is well described by a 3D pure diffusion model with infinite boundary conditions. Assuming that the bleach spot *R_bleach_* is small compared to the droplet size *R_drop_*, the droplet can be effectively treated as an infinite medium, where the prebleach fluorescence concentration is reached far from the bleach spot. *C*^∗^(*R_drop_*, *t*) = 1. This assumption is generally met when *R_drop_*/*R_bleach_* ≥ 2.3 ^113^. Under this condition, the time evolution of the normalized concentration after photobleaching for *r* < *R_bleach_*, satisfying the diffusion equation 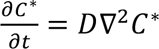, can be written as

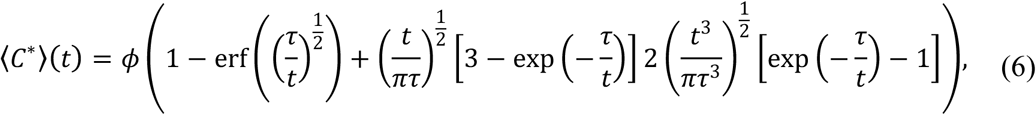

where *φ* is the fraction and *τ* being the recovery time *τ* = *R*^2^_*bleach*_ /*D*, with D being the effective diffusion coefficient. Therefore, fitting the above equation to the experimental concentration profile allows the estimation of *D* and *φ*, as the best-fitting parameters.

### Droplet coalescence analysis

During the 2-hours equilibration phase described in the previous paragraph, droplet formation, sedimentation on the bottom surface and fusion events between touching droplets were monitored by acquiring time-lapse images with a frame rate of 1 fps of a large field of view (image size 222 µm (2048×2048 *pixels*)) using the same microscope, camera and objective described in previous section. For each experiment, about N=50 distinct fusion events were identified by eye and, for each of them, a sub-stack centered on the fusing droplets and starting a few frames before they come in contact was generated.

The contour of the fusing droplets is obtained by applying a Wiener filter of standard deviation 0.2 µm and then a threshold to the images. Upon contact, the aspect ratio *AR* of the fusing droplet was monitored over time. An exponential model was fitted to the obtained *AR*(*t*)

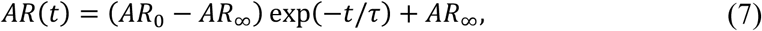

where *AR*_∞_ represents the initial aspect ratio at time *t* = 0 when the fusion process begins, *AR*_∞_ is the asymptotic aspect ratio (i.e. the estimated value of the aspect ratio for long times) and *τ* is a characteristic fusion time. If the fusing droplets are purely viscous, with constant viscosity *η_b_*, one would expect *AR*_∞_ = 1, while *τ* is given by

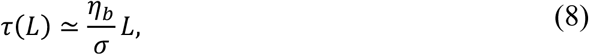

where *σ* is the interfacial tension and 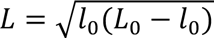 is the characteristic droplet size estimated as the average droplet diameter at the time of contact *t* = 0 *s* − where *L*_0_ and *l*_0_ are the major and minor axis dimensions, respectively. The above relation enables estimating the coalescence velocity 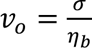 for each coalescence event. In the presence of a non-negligible elastic modulus *E*, the final shape can deviate from perfect sphere leading to a non-vanishing value of *AR*_∞_^46^

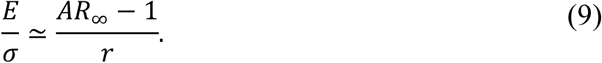

### Cell culture

Cells were grown under standard tissue culture conditions (37 °C, 5% CO_2_). HeLa cells were cultured in Dulbecco’s Modified Eagle’s Medium (DMEM) high glucose (ECB7501L, Euroclone), supplemented with 10% Fetal Bovine Serum (FBS), and 2 mM L-glutamine. BJ cells infected with mCherry-BP1-2 pLPC-Puro plasmid (19835, Addgene) were previously described^14^, and cultured in Minimum Essential Medium (MEM, ECB2071L, Euroclone) supplemented with 10% FBS, 2 mM L-glutamine, 1% non-essential amino acids, and 1% sodium pyruvate. SV40LT-immortalized *Trf2^F/F^ 53bp1^-/-^* Mouse Embryonic Fibroblasts (MEFs) were previously described^54^, and cultured in DMEM supplemented with 15% FBS, 1% non-essential amino acids, 1% L-glutamine, 1% penicillin/streptomycin, and 50 μM β-mercaptoethanol.

### Cell transfection and transduction

Retroviral particles were obtained after transfection of Phoenix-ECO cells (ATCC, CRL-3214), using Lipofection (jetOPTIMUS DNA Transfection Reagent, Polyplus) (for 53BP1-expressing with pLPC) or CaPO_4_ precipitation (for Hit&Run Cre) using 20 µg of retroviral vector plasmid. The viral supernatant was filtered through a 0.45 μm filter, supplemented with 4 μg/mL polybrene, and used to transduce the target cells. For *53bp1* complementation, MEFs were transduced five times over 3 days (8-16 h intervals) and selected for 2 days in 4 µM puromycin for 53BP1 complementation. For Hit&Run Cre, MEFs were transduced six times over 2 days (6-12 h intervals) and harvested at the indicated time, with time point 0 set at 12 h after the first infection.

### Extraction of total RNA

Pelleted HeLa cells were lysed with QIAzol Lysis Reagent (1023537, Qiagen) and RNA was extracted using miRNeasy Mini Kit (217004, Qiagen) according to the manufacturer’s protocol. On-column DNase digestion was performed during the extraction procedure.

### Extraction of genomic DNA

Genomic DNA was extracted from pelleted HeLa cells using DNeasy Blood & Tissue Kit (69506, Qiagen) following the manufacturer’s instructions.

### Immunoblot

Cells were harvested via trypsinization, washed in PBS and lysed in 2× Laemmli buffer. After denaturation for 10 min at 95°C, chromatin shearing was performed with an insulin needle. The lysate was resolved via SDS-PAGE and then transferred onto nitrocellulose membrane (Cytiva). The membrane was blocked in 5% milk in PBS containing 0.1% (v/v) Tween-20 (PBS-T), and immunoblotted using the following antibodies: 53BP1 (ab175933, Abcam), TRF2 (13136, Cell Signaling), β-actin (3700, Cell Signaling), followed by goat anti-rabbit (31460, Invitrogen) or anti-mouse (31430, Invitrogen) IgG-HRP secondary antibodies. Signals were detected according to the manufacturer’s instructions using chemiluminescent detection reagents (Merck) on ChemiDoc (Bio-Rad) imaging system.

### Fluorescence recovery after photobleaching (FRAP) in live cells

BJ cells infected with mCherry-BP1-2 pLPC-Puro plasmid (19835, Addgene), already used in the lab^14^, were seeded on Mattek dishes, and irradiated with 20 Gy using a CellRad irradiator (Faxitron). FRAP was performed at 2 h, 8 h, 24 h, 1 week after irradiation (on different plates for each time point), using a Spinning Disk Olympus (CSU) microscope. Baseline fluorescence was recorded for 5 frames, then bleaching was performed on the complete focus, and recovery was measured for 3 min (1 frame/sec). Bleaching areas were drawn manually with a circle tool. Analysis was performed with a custom Fiji^109^ plugin. Briefly, the script first performs an automated segmentation to identify individual nuclei. For each of the detected nuclei, a cropped movie enclosing it is generated and the Image-J plugin *Stack-Reg* for rigid image registration is executed to compensate for cell small movements. The fluorescence recovery ROI area is then detected by comparing the pre-bleach and the bleach frame. For each nucleus, the mean fluorescence intensity is measured both within the bleached ROI and across the entire nucleus over the full movie duration. To correct for photobleaching, the ROI data are normalized to the fluorescence signal of the whole nucleus. The post-bleach fluorescence recovery data are then scaled to range from 0 to 1 and fitted with a single exponential recovery equation (y=1−exp(−a∗x)). This fitting process allows for the calculation of key kinetic parameters, including the half-time of recovery (*t_1/2_*), which provides a quantitative measure of protein turnover and internal mobility within the foci.

### Fluorescence in situ hybridization (FISH)

For telomere FISH, MEFs were treated with 0.2 μg/mL Colcemid in the last 1-2 h before collection by trypsinization. Harvested cells were swollen in a hypotonic solution of 0.055-0.075 M KCl at 37°C for 15-30 min before fixation in methanol/acetic acid (3:1) overnight at 4°C. Cells were dropped onto glass slides and allowed to age overnight before dehydration through an ethanol series of 70%, 95% and 100%. Telomere ends were hybridized with Alexa Fluor 488-OO-(TTAGGG)_3_ in hybridization solution [70% formamide, 1 mg/mL blocking reagent (11096176001, Roche), and 10 mM Tris-HCl pH 7.2] overnight at 4°C followed by two washes in 70% formamide, 0.1% Bovine Serum Albumin (BSA), 10 mM Tris-HCl pH 7.2 for 15 min each, and three washes in PBS for 5 min each. Chromosomal DNA was counterstained with the addition of 4′,6-diamidino-2-phenylindole (DAPI) (D1306, Invitrogen) to the second wash. Slides were dehydrated through an ethanol series of 70%, 95% and 100%, left to air-dry and mounted in antifade reagent (Prolong Gold Antifade, P36934, Invitrogen). Images were acquired on a DeltaVision RT microscope system (Applied Precision) with a PlanApo 60 × 1.40 NA objective lens (Olympus America, Inc.) at 1 × 1 binning and multiple 0.2 μm Z-stacks using SoftWoRx software and deconvolved. 2D-maximum intensity projection images were obtained using SoftWoRx software. Chromosome-type fusions were analyzed using Fiji software^109^.

### Statistics and reproducibility

The number *n* of times experiments were repeated with similar results and the statistic tests used to calculate significance are provided in the figure legend for each experiment. GraphPad Prism 10 and MATLAB were used to generate graphs and to perform statistical analysis. *P* values of statistical significance are indicated as n.s. *P* > 0.05; * *P* < 0.05; ** *P* < 0.01; *** *P*< 0.001; **** *P* < 0.0001.

## DATA AVAILABILITY

NMR chemical shift assignment of 53BP1 F6 is in the process of deposition to BMRB (BMRB ID TBD). Plasmid for the pTHMT 53BP1 F6 is in the process of deposition to Addgene and will be available from https://www.addgene.org/Nicolas_Fawzi/.

## ACKNOWLEDGEMENTS

We would like to thank: A. De Antoni (IFOM, Milan, Italy) and A. de Marco (Nova Gorica University, Nova Gorica, Slovenia) for providing plasmids; G. Ciossani and L. Scietti (IEO, Milan, Italy) for support with FP studies and useful suggestions about protein purification, chromatography techniques, and AlphaFold 3 predictions; P. Bonaiuti (IFOM, Milan, Italy) for helpful advice on droplets analysis and quantification; S. Barozzi and D. Parazzoli (IFOM, Advanced Light Microscopy Core Facility – RRID:SCR_026866, Milan, Italy) for support with imaging experiments; E. Maspero and S. Polo (IFOM, Milan, Italy) for useful suggestions about protein purification and chromatography techniques; E. Guccione (Mount Sinai Hospital, New York, USA) for useful suggestions about peptides synthesis; Tilo Schorn and Maura Francolini (University of Milan, Italy) for support with FRAP experiments; Giuliano Zanchetta (University of Milan, Italy) for useful discussion about biophysical characterization of biomolecular condensates.

## AUTHOR CONTRIBUTIONS

**F.T.** wrote the manuscript, analyzed the experiments (except for computational, NMR, biophysical and telomere fusions analyses), and assembled the manuscript figures; expressed and purified 53BP1 protein fragments; performed phase separation assays, FRAP in live cells, RNA extraction; cloned plasmids expressing 53BP1 WT, R→A and R→K mutants for retroviral infection in MEFs; performed AlphaFold 3 predictions; provided help with biophysical experiments and recombinant protein quality control analyses. **M.C.** wrote the manuscript, and assembled part of the manuscript figures; designed, cloned, expressed and purified 53BP1 protein fragments; conceived, performed and analyzed phase separation assays and FP experiments; provided help in setting-up and performing DDM experiments; performed RNA extraction. **O.S.** purified 53BP1 F6 fragment and cloned and purified its R→K form; performed and analyzed phase separation assays and EMSAs; performed DNA and RNA extractions; performed AlphaFold 3 predictions. **T.Z.** and **S.Cu.** subcloned and purified 53BP1 F6 fragment for NMR experiments; conceived, performed and analyzed NMR experiments. **M.B.** performed and analyzed the biophysical experiments, and **S.Co.** provided help with the experimental setup. **A.d.S.** performed and analyzed the telomere fusion experiments in MEFs. **G.O.** purified 53BP1 protein fragments and PS protease, and provided helpful suggestions about protein purification and chromatographic techniques. **F.O.** and **S.M.** conceived and designed the Fiji plugins to quantify liquid droplets in Differential Interference Contrast (DIC) microscopy, and to analyze FRAP in cells. **A.G.** performed the quality control analyses on the purified recombinant proteins. **A.M.** supervised A.G. and edited the manuscript. **M.F.** edited the manuscript; conceived and performed computational analyses on 53BP1 protein fragments. **N.L.F.** supervised T.Z. and S.Cu. and edited the manuscript; conceived and coordinated NMR experiments. **F.L.** supervised A.d.S. and edited the manuscript; conceived and coordinated telomere fusion experiments. **F.G.** supervised M.B. and S.Co. and edited the manuscript; conceived the biophysical experiments, and provided help with droplets quantification and statistical analysis. **F.d.A.d.F.** conceived the study, coordinated and supervised the work, procured funding, and revised the manuscript. All authors reviewed and approved the manuscript.

## DECLARATION OF GENERATIVE AI AND AI-ASSISTED TECHNOLOGIES IN THE WRITING PROCESS

During the preparation of this work the authors used ChatGPT and Gemini to improve language and readability. After using this tool, the authors reviewed and edited the content as needed and take full responsibility for the content of the publication.

## EXTENDED DATA

**Extended Data Fig. 1:**
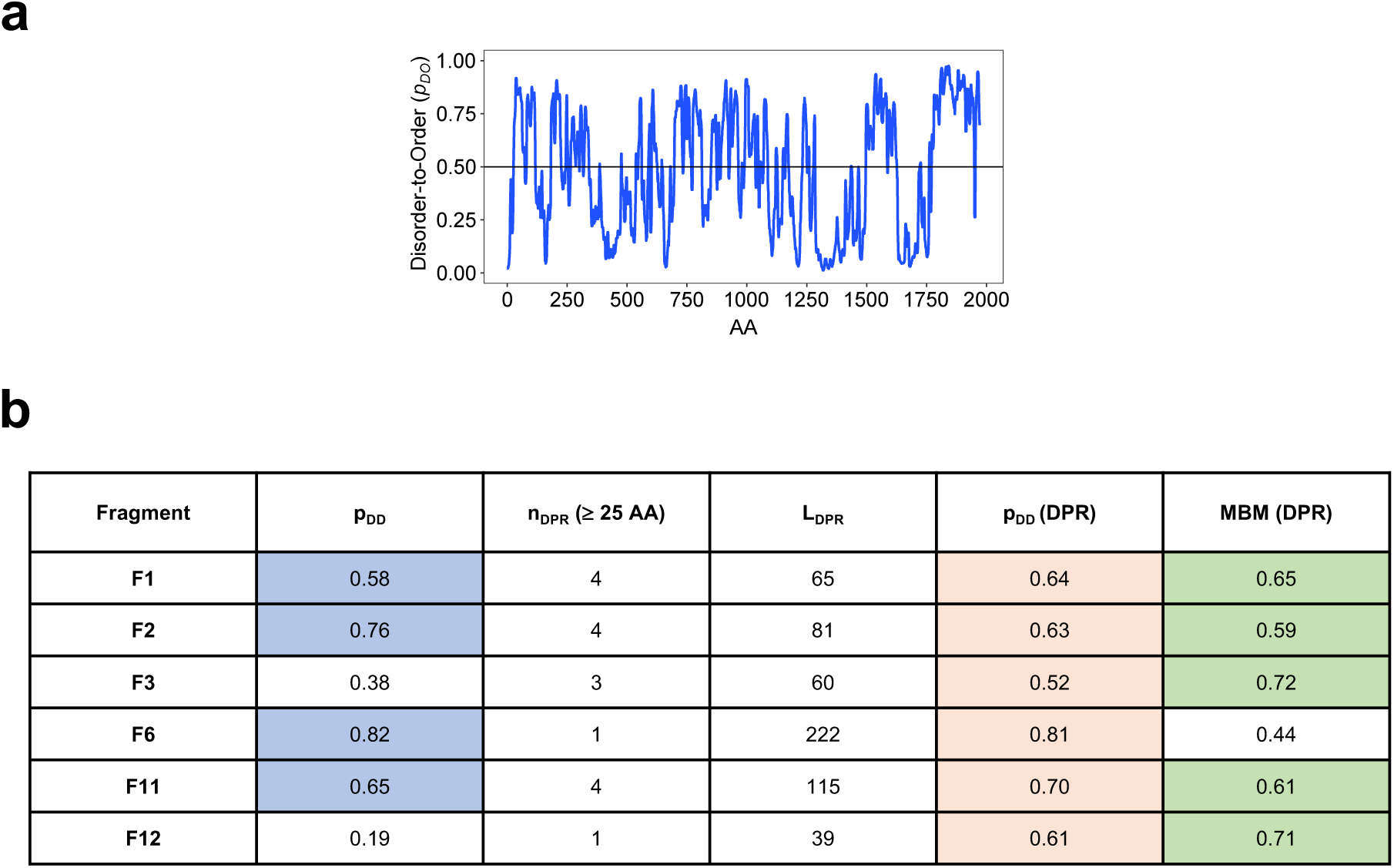
Predicted disordered interaction properties and LLPS propensity of 53BP1 and its fragments. **(a)** Graph plotting the probability of disorder-to-order transitions upon binding (p_DO_) for full-length 53BP1, as computed by FuzPred. Scores above 0.5 (threshold marked by the horizontal black line) indicate transitions to ordered states upon binding. **(b)** Summary table of FuzPred and FuzDrop results for 53BP1 fragments. Colored cells indicate values exceeding the defined threshold for each parameter. The probability to form disordered interactions (p_DD_) is defined as the median of the residue-based values of the fragment as computed by the FuzPred method^25^. Fragments with dominantly disordered interactions (p_DD_ > 0.5) are colored in blue. DPRs were defined as stretches of at least 25 consecutive residues with p_DP_ ≥ 0.60 as computed by the FuzDrop method^27^. The number (n_DPR_) and the average length of the droplet-promoting regions (L_DPR_) are displayed. F1-F3 are equipped with multiple, shorter droplet-promoting regions, while F11 with multiple, longer DPRs. All DPRs exhibit disordered interactions (p_DD_ (DPR) > 0.5). The MBM was used to assess the propensity of DPRs to convert into a more solid-like state^81^. DPRs defined as context-dependent (MBM ≥ 0.59) are colored in green.

**Extended Data Fig. 2:**
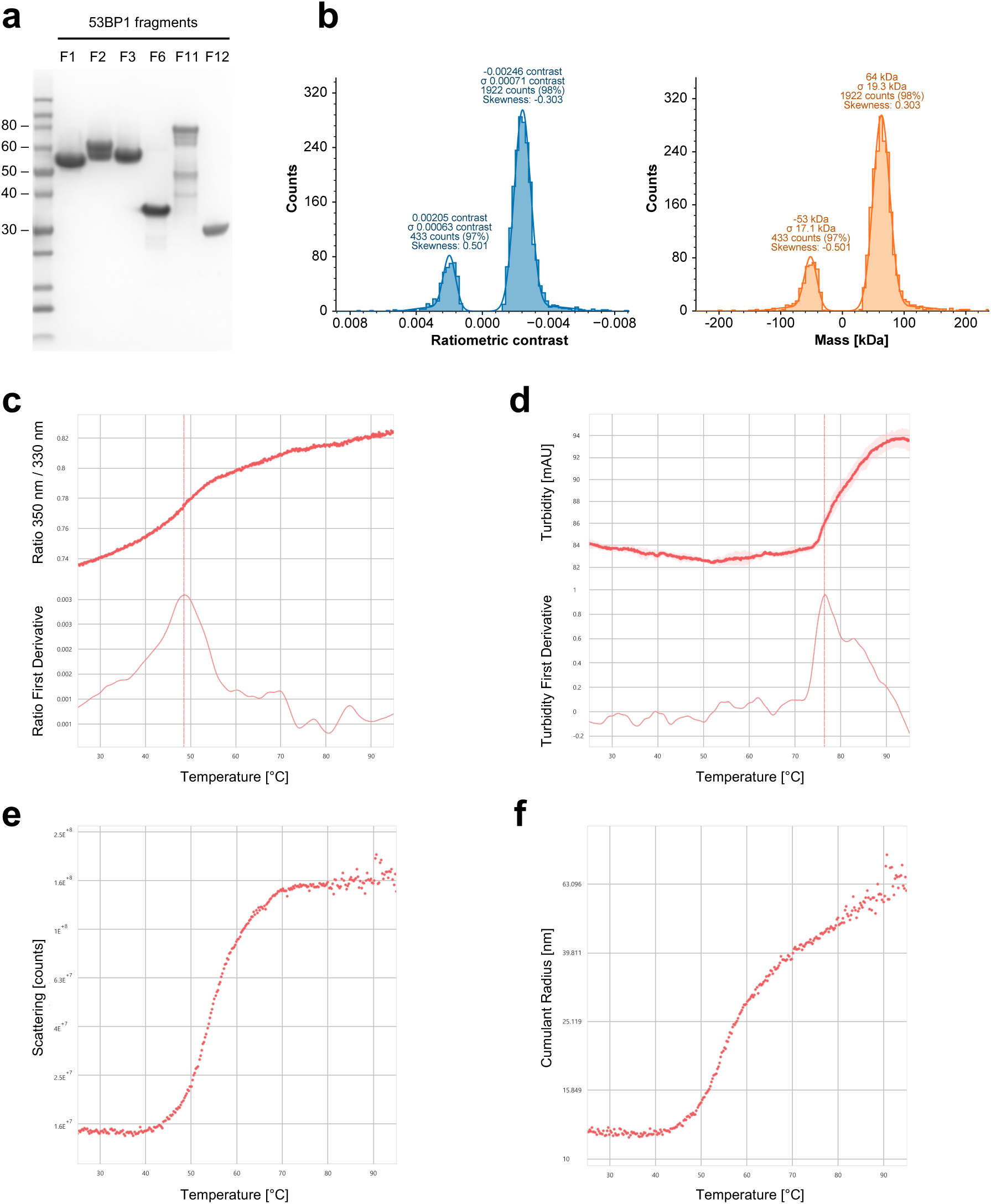
Protein purification of recombinant 53BP1 fragments and quality control analyses. **(a)** SDS-PAGE gel (Coomassie-stained) showing all the purified recombinant 53BP1 fragments used in this study, at their highest achievable purity. **(b)** Mass photometry analysis of purified 53BP1 F6 fragment confirms sample monodispersity and homogeneity. Raw data acquisition at 200 nM displayed on a ratiometric contrast scale (left). Mass distribution at 200 nM obtained after calibration using BSA and BAM standards (right). Measurements were performed in SEC buffer using a TwoMP Instrument (Refeyn). Each condition was tested in duplicate. **(c-f)** Thermal unfolding analysis of purified 53BP1 F6 fragment confirms protein stability. **(c)** nanoDSF F_350/330nm_ profile (top) and its first derivative (bottom). T_m_ = 48.57 ± 0.76°C (dashed line). **(d)** Turbidity (backreflection) profile (top) and its first derivative (bottom). T_agg_ = 76.48 ± 0.07 °C (dashed line). **(e)** Scattering profile (SLS). T_onset_ = 43.50 ± 0.51 °C. **(f)** Cumulant Radius analysis of DLS data. T_onset_ = 47.01 ± 0.24 °C. All the experiments were performed in triplicate at 2 mg/mL in SEC buffer, using a 25 to 95 °C temperature gradient with a ramp of 0.5 °C/min, on a Prometheus Panta NT.48 Instrument (NanoTemper GmBH).

**Extended Data Fig. 3:**
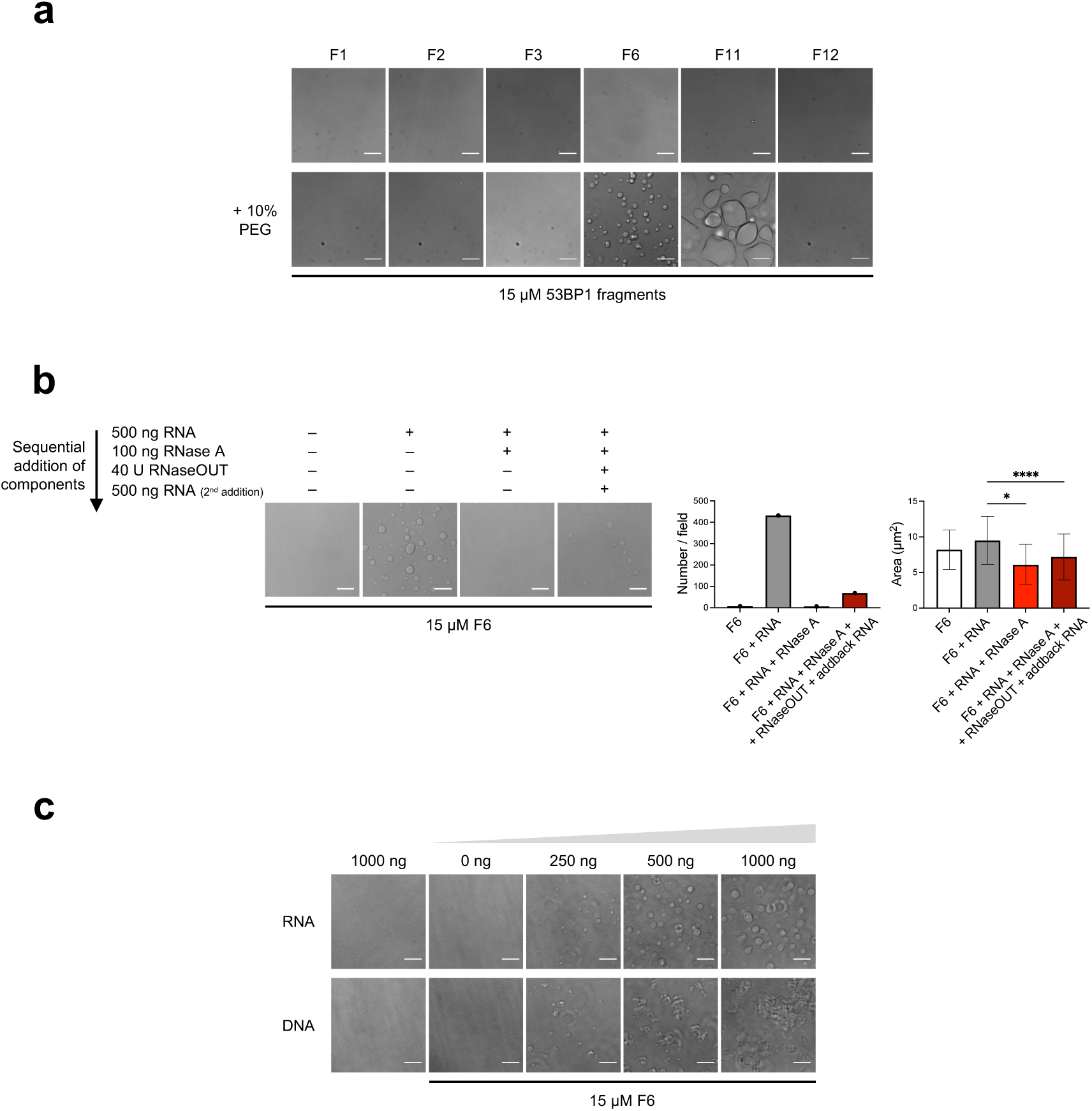
53BP1 OD and its C-terminal disordered region are sufficient for RNA-mediated LLPS *in vitro*. **(a)** Representative images of 53BP1 fragment droplets formation in presence or absence of 10% PEG. The experiment was performed in duplicate, with similar results. DIC channel is shown; scale bar = 10 µm. **(b)** Representative images of F6 droplets dissolution and reformation after RNase A treatment and inhibition, with their relative quantification of droplets number per field and mean droplet area. After droplets formation with RNA, RNase A was added to the well, and imaged 30 min post-treatment; the mixture was incubated with RNaseOUT for 15 min to inactivate RNase A, followed by addition of extra RNA and imaging 30 min later. Data are presented as mean ± SD from *n* = 1 independent experiments. Statistical analysis was performed using one-way ANOVA, followed by Dunnett’s post hoc test against “RNA” group (* *P* < 0.05, **** *P* < 0.0001). DIC channel is shown; scale bar = 10 µm. **(c)** Representative images of F6 behavior in presence of increasing amounts of cellular RNA or genomic DNA. The experiment was performed in triplicate, with similar results. DIC channel is shown; scale bar = 10 µm.

**Extended Data Fig. 4:**
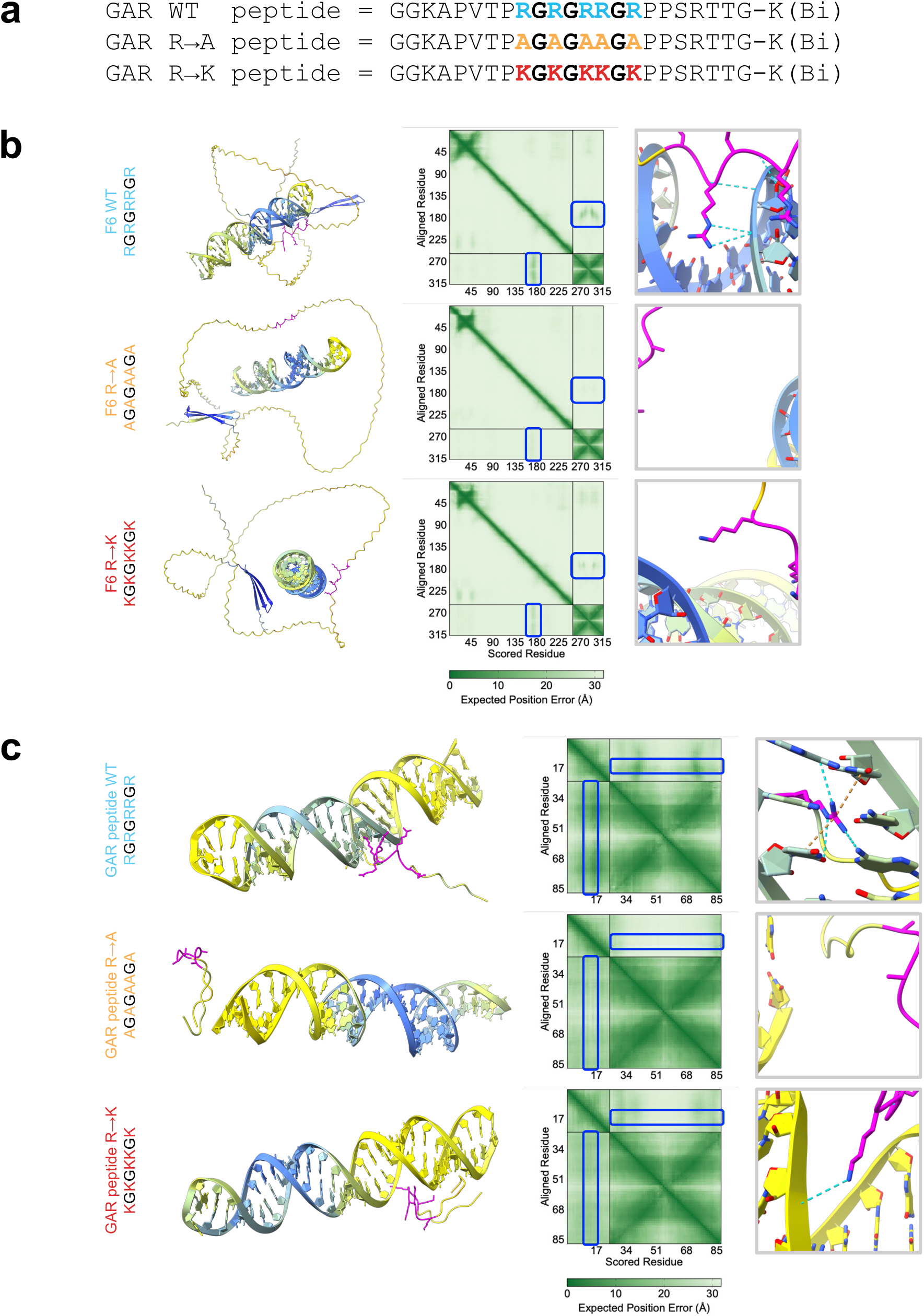
The GAR motif of 53BP1 is predicted to mediate protein-RNA interactions. **(a)** Sequence of the synthetic GAR peptides used in this study, comprising 53BP1 GAR motif in wild-type form with arginines (GAR WT), or mutated to alanines (GAR R→A) or to lysines (GAR R→K). **(b-c)** Representative images of AlphaFold 3-predicted structure of WT, R→A and R→K F6 fragment **(b)** or GAR peptides **(c)** with 63-nt synthetic ssRNA. The stoichiometric ratio of protein or peptide to RNA is 1:1. The predictions were run 10 times for each condition with 10 autogenerated seeds, yielding similar results. The structure is colored by pLDDT. The GAR motif is colored magenta. In the middle, the predicted aligned error (PAE) plot. Black lines delineate borders between different entities. Blue boxes indicate regions corresponding to the GAR motif. On the right, a zoom-in of the simulated interaction between the GAR motif and RNA. Dashed lines represent putative hydrogen bonds (cyan), or cation-π interactions (orange).

**Extended Data Fig. 5:**
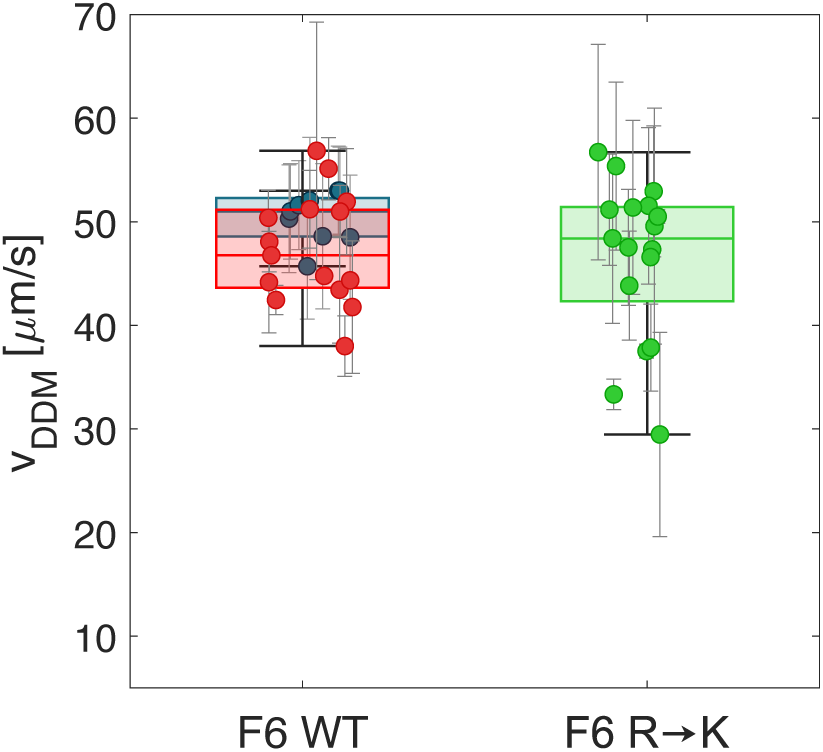
Biophysical properties of F6 WT and F6 R→K mutant condensates. Capillary velocity obtained with DDM on both F6 WT and F6 R→K mutant droplets. Blue dots correspond to large F6 WT droplets (with a diameter larger that 7 µm), for which the capillary velocity is obtained by fitting a quadratic function to the *q*-dependent relaxation rate *Γ*(*q*), as described in main text. Red and green dots correspond to small F6 WT and F6 R→K mutant droplets, respectively. In these cases, the number of available *q*-values is not large enough to perform a robust fitting procedure and the capillary velocity is alternatively estimated as 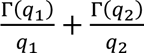, where 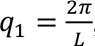, 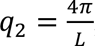, and *L* ≅ 4 *μm* is the size of the considered simplified procedure is confirmed by the good agreement between the results obtained from small and large F6 WT droplets. No statistically significant difference is observed between the capillary velocity on the surface of F6 WT and F6 R→K mutant droplets. All the observations have been performed >2 days after sample preparation. Data from the large F6 WT droplets condition (blue dots) are the same as shown in Fig. 5c to facilitate direct comparison with additional conditions. Data are presented as box plots. Statistical analysis was performed using Mann-Whitney U-test or Wilcoxon rank sum test.

## REFERENCES

1. Hyman, A. A., Weber, C. A. & Jülicher, F. Liquid-Liquid Phase Separation in Biology. Annu. Rev. Cell Dev. Biol. 30, 39–58 (2014).

2. Banani, S. F., Lee, H. O., Hyman, A. A. & Rosen, M. K. Biomolecular condensates: organizers of cellular biochemistry. Nat. Rev. Mol. Cell Biol. 18, 285–298 (2017).

3. Boeynaems, S. et al. Protein Phase Separation: A New Phase in Cell Biology. Trends Cell Biol. 28, 420–435 (2018).

4. Shin, Y. & Brangwynne, C. P. Liquid phase condensation in cell physiology and disease. Science (80-. ). 357, eaaf4382 (2017).

5. Alberti, S. & Hyman, A. A. Biomolecular condensates at the nexus of cellular stress, protein aggregation disease and ageing. Nat. Rev. Mol. Cell Biol. 22, 196–213 (2021).

6. Wang, B. et al. Liquid–liquid phase separation in human health and diseases. Signal Transduct. Target. Ther. 6, 290 (2021).

7. Mehta, S. & Zhang, J. Liquid–liquid phase separation drives cellular function and dysfunction in cancer. Nat. Rev. Cancer 22, 239–252 (2022).

8. Roden, C. & Gladfelter, A. S. RNA contributions to the form and function of biomolecular condensates. Nat. Rev. Mol. Cell Biol. 22, 183–195 (2021).

9. Han, T. W., Portz, B., Young, R. A., Boija, A. & Klein, I. A. RNA and condensates: Disease implications and therapeutic opportunities. Cell Chem. Biol. 31, 1593–1609 (2024).

10. d’Adda di Fagagna, F. Living on a break: cellular senescence as a DNA-damage response. Nat. Rev. Cancer 8, 512–522 (2008).

11. Jackson, S. P. & Bartek, J. The DNA-damage response in human biology and disease. Nature 461, 1071–1078 (2009).

12. Jeggo, P. A. & Löbrich, M. DNA double-strand breaks: their cellular and clinical impact? Oncogene 26, 7717–7719 (2007).

13. Michelini, F. et al. From “Cellular” RNA to “Smart” RNA: Multiple Roles of RNA in Genome Stability and Beyond. Chem. Rev. 118, 4365–4403 (2018).

14. Michelini, F. et al. Damage-induced lncRNAs control the DNA damage response through interaction with DDRNAs at individual double-strand breaks. Nat. Cell Biol. 19, 1400–1411 (2017).

15. Burger, K., Schlackow, M., Gullerova, M. & Yp, C. T. D. Tyrosine kinase c-Abl couples RNA polymerase II transcription to DNA double-strand breaks. Nucleic Acids Res. 47, 3467–3484 (2019).

16. Francia, S. et al. Site-specific DICER and DROSHA RNA products control the DNA-damage response. Nature 488, 231–235 (2012).

17. Burger, K. et al. Nuclear phosphorylated Dicer processes double-stranded RNA in response to DNA damage. J. Cell Biol. 216, 2373–2389 (2017).

18. Francia, S., Cabrini, M., Matti, V., Oldani, A. & d’Adda di Fagagna, F. DICER, DROSHA and DNA damage response RNAs are necessary for the secondary recruitment of DNA damage response factors. J. Cell Sci. 129, 1468–1476 (2016).

19. Mirman, Z. & de Lange, T. 53BP1: a DSB escort. Genes Dev. 34, 7–23 (2020).

20. Pessina, F. et al. Functional transcription promoters at DNA double-strand breaks mediate RNA-driven phase separation of damage-response factors. Nat. Cell Biol. 21, 1286–1299 (2019).

21. Kilic, S. et al. Phase separation of 53BP1 determines liquid-like behavior of DNA repair compartments. EMBO J. 38, e101379 (2019).

22. Kelliher, J. L. et al. Evolved histone tail regulates 53BP1 recruitment at damaged chromatin. Nat. Commun. 15, 4634 (2024).

23. Mészáros, B., Erdős, G. & Dosztányi, Z. IUPred2A: context-dependent prediction of protein disorder as a function of redox state and protein binding. Nucleic Acids Res. 46, W329–W337 (2018).

24. Fuxreiter, M. & Vendruscolo, M. Generic nature of the condensed states of proteins. Nat. Cell Biol. 23, 587–594 (2021).

25. Miskei, M., Horvath, A., Vendruscolo, M. & Fuxreiter, M. Sequence-Based Prediction of Fuzzy Protein Interactions. J. Mol. Biol. 432, 2289–2303 (2020).

26. Iwabuchi, K. et al. Potential Role for 53BP1 in DNA End-joining Repair through Direct Interaction with DNA *. J. Biol. Chem. 278, 36487–36495 (2003).

27. Hardenberg, M., Horvath, A., Ambrus, V., Fuxreiter, M. & Vendruscolo, M. Widespread occurrence of the droplet state of proteins in the human proteome. Proc. Natl. Acad. Sci. 117, 33254–33262 (2020).

28. Vendruscolo, M. & Fuxreiter, M. Sequence Determinants of the Aggregation of Proteins Within Condensates Generated by Liquid-liquid Phase Separation. J. Mol. Biol. 434, 167201 (2022).

29. McGuffin, L. J., Bryson, K. & Jones, D. T. The PSIPRED protein structure prediction server. Bioinformatics 16, 404–405 (2000).

30. Jain, A. & Vale, R. D. RNA phase transitions in repeat expansion disorders. Nature 546, 243–247 (2017).

31. Kroschwald, S., Maharana, S. & Simon, A. Hexanediol: a chemical probe to investigate the material properties of membrane-less compartments. Matters 1–6 (2017) doi:10.19185/matters.201702000010.

32. Zhang, L. et al. 53BP1 regulates heterochromatin through liquid phase separation. Nat. Commun. 13, 360 (2022).

33. Zheng, T. et al. Molecular insights into the effect of 1,6-hexanediol on FUS phase separation. EMBO J. 44, 2725–2740 (2025).

34. Leung, A. K. L. L. Poly(ADP-ribose): A Dynamic Trigger for Biomolecular Condensate Formation. Trends Cell Biol. 30, 370–383 (2020).

35. Alemasova, E. E. & Lavrik, O. I. A sePARate phase? Poly(ADP-ribose) versus RNA in the organization of biomolecular condensates. Nucleic Acids Res. 50, 10817–10838 (2022).

36. Shakya, A. & King, J. T. DNA Local-Flexibility-Dependent Assembly of Phase-Separated Liquid Droplets. Biophys. J. 115, 1840–1847 (2018).

37. Muzzopappa, F., Hertzog, M. & Erdel, F. DNA length tunes the fluidity of DNA-based condensates. Biophys. J. 120, 1288–1300 (2021).

38. Howe, J. et al. LC8 enhances 53BP1 foci through heterogeneous bridging of 53BP1 oligomers. Elife 14, e102179 (2025).

39. Marsh, J. A., Singh, V. K., Jia, Z. & Forman-Kay, J. D. Sensitivity of secondary structure propensities to sequence differences between α- and γ-synuclein: Implications for fibrillation. Protein Sci. 15, 2795–2804 (2006).

40. Thandapani, P., O’Connor, T. R., Bailey, T. L. L. L. & Richard, S. Defining the RGG/RG Motif. Mol. Cell 50, 613–623 (2013).

41. Chong, P. A., Vernon, R. M. & Forman-Kay, J. D. RGG/RG Motif Regions in RNA Binding and Phase Separation. J. Mol. Biol. 430, 4650–4665 (2018).

42. Boisvert, F.-M., Alexandre, R., Stéphane, R. & and Doherty, A. J. The GAR Motif of 53BP1 is Arginine Methylated by PRMT1 and is Necessary for 53BP1 DNA Binding Activity. Cell Cycle 4, 1834–1841 (2005).

43. Adams, M. M. et al. 53BP1 Oligomerization is Independent of its Methylation by PRMT1. Cell Cycle 4, 1854–1861 (2005).

44. Leriche, M. et al. 53BP1 interacts with the RNA primer from Okazaki fragments to support their processing during unperturbed DNA replication. Cell Rep. 42, (2023).

45. Abramson, J. et al. Accurate structure prediction of biomolecular interactions with AlphaFold 3. Nature 630, 493–500 (2024).

46. Aarts, D. G. A. L., Schmidt, M. & Lekkerkerker, H. N. W. Direct Visual Observation of Thermal Capillary Waves. Science (80-. ). 304, 847–850 (2004).

47. Cerbino, R. & Trappe, V. Differential Dynamic Microscopy: Probing Wave Vector Dependent Dynamics with a Microscope. Phys. Rev. Lett. 100, 188102 (2008).

48. Giavazzi, F., Brogioli, D., Trappe, V., Bellini, T. & Cerbino, R. Scattering information obtained by optical microscopy: Differential dynamic microscopy and beyond. Phys. Rev. E 80, 31403 (2009).

49. Wang, J. & McGorty, R. Measuring capillary wave dynamics using differential dynamic microscopy. Soft Matter 15, 7412–7419 (2019).

50. Pawar, A. B., Caggioni, M., Hartel, R. W. & Spicer, P. T. Arrested coalescence of viscoelastic droplets with internal microstructure. Faraday Discuss. 158, 341–350 (2012).

51. Narhe, R., Beysens, D. & Nikolayev, V. S. Dynamics of Drop Coalescence on a Surface: The Role of Initial Conditions and Surface Properties. Int. J. Thermophys. 26, 1743–1757 (2005).

52. Dimitrova, N., Chen, Y.-C. C. M. C. M., Spector, D. L. & De Lange, T. 53BP1 promotes non-homologous end joining of telomeres by increasing chromatin mobility. Nature 456, 524–528 (2008).

53. Lottersberger, F., Bothmer, A., Robbiani, D. F., Nussenzweig, M. C. & de Lange, T. Role of 53BP1 oligomerization in regulating double-strand break repair. Proc. Natl. Acad. Sci. 110, 2146–2151 (2013).

54. Lottersberger, F., Karssemeijer, R. A. A., Dimitrova, N., de Lange, T. & de Lange, T. 53BP1 and the LINC Complex Promote Microtubule-Dependent DSB Mobility and DNA Repair. Cell 163, 880–893 (2015).

55. Silver, D. P. & Livingston, D. M. Self-Excising Retroviral Vectors Encoding the Cre Recombinase Overcome Cre-Mediated Cellular Toxicity. Mol. Cell 8, 233–243 (2001).

56. Pryde, F. et al. 53BP1 exchanges slowly at the sites of DNA damage and appears to require RNA for its association with chromatin. J. Cell Sci. 118, 2043–2055 (2005).

57. Botuyan, M. V. et al. Structural Basis for the Methylation State-Specific Recognition of Histone H4-K20 by 53BP1 and Crb2 in DNA Repair. Cell 127, 1361–1373 (2006).

58. Ketley, R. F. et al. DNA double-strand break-derived RNA drives TIRR/53BP1 complex dissociation. Cell Rep. 41, (2022).

59. Charier, G. et al. The Tudor Tandem of 53BP1: A New Structural Motif Involved in DNA and RG-Rich Peptide Binding. Structure 12, 1551–1562 (2004).

60. Gallivan, J. P. & Dougherty, D. A. Cation-π interactions in structural biology. Proc. Natl. Acad. Sci. 96, 9459–9464 (1999).

61. Wang, J. et al. A Molecular Grammar Governing the Driving Forces for Phase Separation of Prion-like RNA Binding Proteins. Cell 174, 688–699.e16 (2018).

62. Kappel, K. et al. Characterizing protein sequence determinants of nuclear condensates by high-throughput pooled imaging with CondenSeq. Nat. Methods 22, 1464–1475 (2025).

63. Murthy, A. C. et al. Molecular interactions contributing to FUS SYGQ LC-RGG phase separation and co-partitioning with RNA polymerase II heptads. Nat. Struct. Mol. Biol. 28, 923–935 (2021).

64. Nomoto, A., Nishinami, S. & Shiraki, K. Solubility Parameters of Amino Acids on Liquid–Liquid Phase Separation and Aggregation of Proteins. Front. Cell Dev. Biol. **Volume** 9-, (2021).

65. Zheng, T. et al. RNA modulates FUS condensate assembly, dynamics, and aggregation through diverse molecular contacts. bioRxiv 2025.12.13.694118 (2025) doi:10.64898/2025.12.13.694118.

66. Puterbaugh, R. Z. et al. The Structural Basis for RNA Binding and Recognition of the Disordered Prion-Like Domain of TDP-43. *bioRxiv* 2025.12.22.695782 (2025) doi:10.64898/2025.12.22.695782.

67. Rai, R. et al. The function of classical and alternative non-homologous end-joining pathways in the fusion of dysfunctional telomeres. EMBO J. 29, 2598–2610 (2010).

68. Fumagalli, M. et al. Telomeric DNA damage is irreparable and causes persistent DNA-damage-response activation. Nat. Cell Biol. 14, 355–365 (2012).

69. Fumagalli, M., Rossiello, F., Mondello, C. & d’Adda di Fagagna, F. Stable Cellular Senescence Is Associated with Persistent DDR Activation. PLoS One 9, e110969 (2014).

70. Rodier, F. et al. Persistent DNA damage signalling triggers senescence-associated inflammatory cytokine secretion. Nat. Cell Biol. 11, 973–979 (2009).

71. Hewitt, G. et al. Telomeres are favoured targets of a persistent DNA damage response in ageing and stress-induced senescence. Nat. Commun. 3, 708 (2012).

72. Lin, Y., Protter, D. S. W. S. W. S. W., Rosen, M. K. K. K. & Parker, R. Formation and Maturation of Phase-Separated Liquid Droplets by RNA-Binding Proteins. Mol. Cell 60, 208–219 (2015).

73. Molliex, A. et al. Phase Separation by Low Complexity Domains Promotes Stress Granule Assembly and Drives Pathological Fibrillization. Cell 163, 123–133 (2015).

74. Zhang, H. et al. RNA Controls PolyQ Protein Phase Transitions. Mol. Cell 60, 220– 230 (2015).

75. Alberti, S. & Dormann, D. Liquid–Liquid Phase Separation in Disease. Annu. Rev. Genet. 53, 171–194 (2019).

76. Elbaum-Garfinkle, S. Matter over mind: Liquid phase separation and neurodegeneration. J. Biol. Chem. 294, 7160–7168 (2019).

77. Babinchak, W. M. & Surewicz, W. K. Liquid–Liquid Phase Separation and Its Mechanistic Role in Pathological Protein Aggregation. J. Mol. Biol. 432, 1910–1925 (2020).

78. Murakami, T. et al. ALS/FTD Mutation-Induced Phase Transition of FUS Liquid Droplets and Reversible Hydrogels into Irreversible Hydrogels Impairs RNP Granule Function. Neuron 88, 678–690 (2015).

79. Patel, A. et al. A Liquid-to-Solid Phase Transition of the ALS Protein FUS Accelerated by Disease Mutation. Cell 162, 1066–1077 (2015).

80. Maharana, S. et al. RNA buffers the phase separation behavior of prion-like RNA binding proteins. Science (80-. ). 360, 918–921 (2018).

81. Vendruscolo, M. & Fuxreiter, M. Protein condensation diseases: therapeutic opportunities. Nat. Commun. 13, 5550 (2022).

82. Horvath, A., Vendruscolo, M. & Fuxreiter, M. Sequence-based Prediction of the Cellular Toxicity Associated with Amyloid Aggregation within Protein Condensates. Biochemistry 61, 2461–2469 (2022).

83. Hwang, J. W. et al. PRMT5 promotes DNA repair through methylation of 53BP1 and is regulated by Src-mediated phosphorylation. Commun. Biol. 3, 428 (2020).

84. Qamar, S. et al. FUS Phase Separation Is Modulated by a Molecular Chaperone and Methylation of Arginine Cation-&# Interactions. Cell 173, 720–734.e15 (2018).

85. Hofweber, M. et al. Phase Separation of FUS Is Suppressed by Its Nuclear Import Receptor and Arginine Methylation. Cell 173, 706–719.e13 (2018).

86. Ryan, V. H. et al. Mechanistic View of hnRNPA2 Low-Complexity Domain Structure, Interactions, and Phase Separation Altered by Mutation and Arginine Methylation. Mol. Cell 69, 465–479.e7 (2018).

87. Xiang, Y. et al. RNA m6A methylation regulates the ultraviolet-induced DNA damage response. Nature 543, 573–576 (2017).

88. Chen, H. et al. m5C modification of mRNA serves a DNA damage code to promote homologous recombination. Nat. Commun. 11, 2834 (2020).

89. Jimeno, S. et al. ADAR-mediated RNA editing of DNA:RNA hybrids is required for DNA double strand break repair. Nat. Commun. 12, 5512 (2021).

90. Yang, H. et al. The RNA m5C modification in R-loops as an off switch of Alt-NHEJ. Nat. Commun. 14, 6114 (2023).

91. Ries, R. J. et al. m6A enhances the phase separation potential of mRNA. Nature 571, 424–428 (2019).

92. Lee, J.-H. H. et al. Enhancer RNA m6A methylation facilitates transcriptional condensate formation and gene activation. Mol. Cell 81, 3368–3385.e9 (2021).

93. Sun, Y. et al. m1A in CAG repeat RNA binds to TDP-43 and induces neurodegeneration. Nature 623, 580–587 (2023).

94. Zhang, C. et al. METTL3 and N6-Methyladenosine Promote Homologous Recombination-Mediated Repair of DSBs by Modulating DNA-RNA Hybrid Accumulation. Mol. Cell 79, 425–442.e7 (2020).

95. Abakir, A. et al. N6-methyladenosine regulates the stability of RNA:DNA hybrids in human cells. Nat. Genet. 52, 48–55 (2020).

96. Shiromoto, Y., Sakurai, M., Minakuchi, M., Ariyoshi, K. & Nishikura, K. ADAR1 RNA editing enzyme regulates R-loop formation and genome stability at telomeres in cancer cells. Nat. Commun. 12, 1654 (2021).

97. Zhang, B. et al. ADAR1 links R-loop homeostasis to ATR activation in replication stress response. Nucleic Acids Res. 51, 11668–11687 (2023).

98. Mitrea, D. M., Mittasch, M., Gomes, B. F., Klein, I. A. & Murcko, M. A. Modulating biomolecular condensates: a novel approach to drug discovery. Nat. Rev. Drug Discov. 21, 841–862 (2022).

99. Klein, I. A. et al. Partitioning of cancer therapeutics in nuclear condensates. Science (80-. ). 368, 1386–1392 (2020).

100. Rossiello, F. et al. DNA damage response inhibition at dysfunctional telomeres by modulation of telomeric DNA damage response RNAs. Nat. Commun. 8, 13980 (2017).

101. Aguado, J. et al. Inhibition of DNA damage response at telomeres improves the detrimental phenotypes of Hutchinson–Gilford Progeria Syndrome. Nat. Commun. 10, 4990 (2019).

102. Sepe, S. et al. Telomeric DNA damage response mediates neurotoxicity of Aβ42 oligomers in Alzheimer’s disease. EMBO J. 1–34 (2025) 10.1038/s44318-025-00521-1.

103. Rosso, I. et al. Alternative lengthening of telomeres (ALT) cells viability is dependent on C-rich telomeric RNAs. Nat. Commun. 14, 7086 (2023).

104. Hatos, A. et al. FuzPred: a web server for the sequence-based prediction of the context-dependent binding modes of proteins. Nucleic Acids Res. 51, W198–W206 (2023).

105. Meng, E. C. et al. UCSF ChimeraX: Tools for structure building and analysis. Protein Sci. 32, e4792 (2023).

106. Peti, W. & Page, R. Strategies to maximize heterologous protein expression in Escherichia coli with minimal cost. Protein Expr. Purif. 51, 1–10 (2007).

107. Alberti, S. et al. A User’s Guide for Phase Separation Assays with Purified Proteins. J. Mol. Biol. 430, 4806–4820 (2018).

108. Hollandi, R. et al. nucleAIzer: A Parameter-free Deep Learning Framework for Nucleus Segmentation Using Image Style Transfer. Cell Syst. 10, 453–458.e6 (2020).

109. Schindelin, J. et al. Fiji: an open-source platform for biological-image analysis. Nat. Methods 9, 676–682 (2012).

110. Delaglio, F. et al. NMRPipe: A multidimensional spectral processing system based on UNIX pipes. J. Biomol. NMR 6, 277–293 (1995).

111. Skinner, S. P. et al. CcpNmr AnalysisAssign: a flexible platform for integrated NMR analysis. J. Biomol. NMR 66, 111–124 (2016).

112. Castellini, S. et al. Modeling and correction of image drift in dynamic shadowgraphy experiments. Eur. Phys. J. E 47, 25 (2024).

113. Taylor, N. O., Wei, M.-T., Stone, H. A. & Brangwynne, C. P. Quantifying Dynamics in Phase-Separated Condensates Using Fluorescence Recovery after Photobleaching. Biophys. J. 117, 1285–1300 (2019).

