## Supplementary Table 1 for "The glycine-arginine-rich motif of 53BP1 modulates RNA interactions necessary for its liquid-liquid phase separation during DNA Damage Response"

| Primer ID | Sequence (5' → 3') |
|---|---|
| FF1 | CGGGATCCATGGACCCTACTGGAAGTCA |
| Rev361 | CCCTCGAGTTAATCTGAAGAATTCGTGGAA |
| FF302 | CGGGATCCATGACTCAGGAAGACTTGTTTG |
| Rev718 | CCCTCGAGTTAAACTTCCATAGCTTCTGAG |
| FF667 | CGGGATCCATGATCCCTGAGACACCTTG |
| Rev1052 | CCCTCGAGTTAATCCTCACTTCGAGCCTCA |
| FF1231 | CGGGATCCATGCCACATGGCCATGT |
| Rev1483 | CCCTCGAGTTATCCTGGAGAGGAGGC |
| FF1053 | CGGGATCCATGCCCCCCACCA |
| Rev1711 | CCCTCGAGTTATTCACCGGTGTTGTCTC |
| FF1711 | CGGGATCCATGGAACCCTCTGCCCT |
| Rev1972 | CCCTCGAGTTAGTGAGAAACATAATCGTGT |
| FF R(GAR)A | GCAGGGGCAGGGGCAGCAGGCGCACCACCTTCTCGGACC |
| Rev R(GAR)A | TGCGCCTGCTGCCCCTGCCCCTGCAGGCGTGACTGGAGC |
| FF FL_R(GAR)A | GCAGCAGGCGCACCACCTTCTCGGACCACTGG |
| Rev FL_R(GAR)A | CCCTGCCCCTGCAGGCGTGACTGGAGCC |
| FF FL_R(GAR)K | AAGAAGGGCAAGCCACCTTCTCGGACCACTGG |
| Rev FL_R(GAR)K | CCCCTTCCCCTTAGGCGTGACTGGAGCC |

| Synthetic RNA oligo ID | Sequence (5' → 3') |
|---|---|
| 63-nt ssRNA | GGACUCGGACUUGGACUCUGGACUUUGGACUUGGACUUGGACUUCGGACUCGGACUUUGGACU – Biotin |
| CXCR4-FAM | UGUUAGCUGGAGUGAAAACUU – 6-FAM |

| Synthetic peptide ID | Sequence (N-ter → C-ter) |
|---|---|
| GAR WT | GGKAPVTPRGRGRRGRPPSRTTG – K(Biotin) |
| GAR R>A | GGKAPVTPAGAGAAGAPPSRTTG – K(Biotin) |
| GAR R>K | GGKAPVTPKGKGKKGKPPSRTTG – K(Biotin) |

*K(Biotin) = lysine biotinylated on the amino group of its side chain*
